## Supplementary material for "Estimates of the basic reproduction number for rubella using seroprevalence data and indicator-based approaches"

### Supplement

### S1: Summary of the study settings

**Table S1**: Summary of the sources of seroprevalence data from each region for the force of infection estimates for each setting, that is used in this analysis. Force of infection estimates from all of these datasets have been previously used in estimating the global burden of Congenital Rubella Syndrome or in the impact of Measles-Rubella vaccination.

| **Africa** | **Americas** | **Eastern Mediterranean** | **Europe** | **South East Asia** | **Western Pacific** |
|---|---|---|---|---|---|
| Benin, 1993[1] | Argentina (rural), 1967-68[2] | Bahrain, 1981[3] | Czech Republic, <1967[4] | Bangladesh, 2004-05[5] | Australia, <1967[4] |
| Burkina Faso, 2007-8[6] | Argentina (urban), 1967-68[2] | Iran, 1993-95[7] | Denmark, <1967[4] | India (rural Delhi), 1968[8] | Cambodia, 2012[9] |
| Congo, <1991[10] | Argentina (Mar de Plata), 1981[11] | Jordan, 1982-3[12] | Denmark, 1983[13] | India (urban Delhi), 1968[8] | China, 1979-80[14] |
| Cote d'Ivoire, 1975[15] | Brazil, 1967-68[2] | Kuwait, <1978[16] | East Germany, 1990[17] | India (Chandrigarh), 1972-3[8] | Fiji, <1973[18] |
| Cote d'Ivoire, 1985-6[19] | Brazil, 1987[20] | Lebanon, 1980-81[21] | England, <1967[4] | India (Lucknow), 1972-3[8] | Japan (Sapporo), <1967[4] |
| Democratic Republic of the Congo (Kikwit), 2008-9[22] | Brazil (Parana), 1996-8[23] | Morocco, 1969-1970[24] | England, 1986-87[25] | India (Calcutta), 1976[26] | Japan (Ohtsu), <1967[4] |
| Democratic Republic of the Congo (Mikalayi), 2008-9[22] | Canada, <1967[4] | Pakistan, <1997[27] | Finland, 1979[28] | India (Delhi), <1987[29] | Malaysia, <1972[30] |
| Democratic Republic of the Congo (Tshikapa), 2008-9[22] | Chile (rural), 1967-68[2] | Pakistan, 1999-2004[31] | France, <1967[4] | India (Delhi), <1990[32] | Singapore, 1975-79[33] |
| Democratic Republic of the Congo (Vanga), 2008-9[22] | Chile (Santiago), 1967-68[2] | Saudi Arabia, 1989[34] | Kyrgyzstan, 2001[35] | India (rural Vellore), 1999-2000[36] | Central Vietnam, 2009-2010[37] |
| Ethiopia, 1981[38] | Haiti, 2002[39] | Saudi Arabia, 1992-93[40] | Romania, <1989[41] | India (urban Vellore), 1999-2000[36] | |

**Table S1** (continued)

| **Africa** | **Americas** | **Eastern Mediterranean** | **Europe** | **South East Asia** | **Western Pacific** |
|---|---|---|---|---|---|
| Ethiopia (Addis Ababa), 1994[42] | Jamaica (Kingston), 1967-68[2] | Tunisia, <1970[43] | Turkey, 1998[44] | Indonesia, 2007 (*S Reef, personal communication, March 2015*) | |
| Gabon, 1985[45] | Jamaica (rural), 1967-68[2] | Yemen, 1985[46] | Turkey, 2003-04[47] | Nepal, 2008[48] | |
| Ghana, 1997[49] | Mexico, 1987-88[50] | Yemen, 2002-03[51] | Turkey, 2005[52] | Thailand, 1978[53] | |
| Kenya (Eldoret), 2005[54] | Mexico, 1989[55] | | | | |
| Kenya (Kilifi), 1996-9[56, 57] | Panama (Panama City), 1967-68[2] | | | | |
| Madagascar, 1990-1995[58] | Panama (rural), 1967-68[2] | | | | |
| Mozambique, 2002[59] | Peru (Lima), 1967-68[2] | | | | |
| Nigeria, <1978[60] | Peru (rural), 1967-68[2] | | | | |
| Nigeria, <2002[61] | Peru, 2003[62] | | | | |
| Nigeria, 2007-8[63] | Trinidad, 1966-7[64] | | | | |
| Senegal, 1996-2001[65] | Trinidad (rural), 1967-8[2] | | | | |
| South Africa, 2003[66] | Trinidad (Port au Spain), 1967-68[2] | | | | |
| Tanzania (Mwanza), 2012-13[67] | Uruguay (rural) 1967-68[2] | | | | |
| Zambia, 1979-80[68] | Uruguay (urban) 1967-68[2] | | | | |
| | USA (Atlanta), <1967[4] | | | | |
| | USA (Houston), <1967[4] | | | | |

#### S2 Completeness of the indicator data

**Table S2** Number of studies for which the year in which the indicator was available differed by less than 5 years, 5-10 years or more than 10 years from the year in which the data were collected. For studies with an unspecified year of data collection, the difference was calculated as the difference between the publication year and the year for which the indicator was extracted. The right-most column cites the source from which the data was obtained.

| | **Number of years difference between the year in which the indicator was available and the study year** | | | | **Data source** |
|---|---|---|---|---|---|
| | **<5** | **5-10** | **>10** | **Indicator not available** | |
| **Demographic** | | | | | |
| Proportion of the population aged 0-4 | 98 | 0 | 0 | 0 | [69] |
| Proportion of the population aged 0-14 | 98 | 0 | 0 | 0 | [69] |
| Proportion of the population aged 65+ | 98 | 0 | 0 | 0 | [69] |
| Lifetime risk of maternal death (1 in: rate varies by country) | 46 | 9 | 43 | 0 | [70] |
| Probability of dying before age 5 (per 1000 live births) | 98 | 0 | 0 | 0 | [69] |
| Crude death rate per 1000 population | 98 | 0 | 0 | 0 | [69] |
| Life expectancy at birth (both sexes) | 98 | 0 | 0 | 0 | [69] |
| Total fertility rate (live births per woman) | 98 | 0 | 0 | 0 | [69] |
| Mean age of child-bearing | 98 | 0 | 0 | 0 | [69] |
| Population growth rate (Average annual rate of population change (percentage)) | 98 | 0 | 0 | 0 | [69] |
| Population density (people per sq. km of land area) | 98 | 0 | 0 | 0 | [70] |
| Urban population (% of total) | 98 | 0 | 0 | 0 | [70] |
| **Economic** | | | | | |
| Income share held by highest 10% | 36 | 6 | 50 | 6 | [70] |
| Income share held by highest 20% | 36 | 6 | 50 | 6 | [70] |
| Income share held by lowest 10% | 36 | 6 | 50 | 6 | [70] |
| Income share held by lowest 20% | 36 | 6 | 50 | 6 | [70] |
| Poverty gap at $1.90 a day (2011 PPP) (%) | 40 | 11 | 41 | 6 | [70] |
| Poverty gap at $3.20 a day (2011 PPP) (% of population) | 40 | 11 | 41 | 6 | [70] |

**Table S2** (continued)

| | Number of years difference between the year in which the indicator was available and the study year | | | | Data source |
|---|---|---|---|---|---|
| | <5 | 5-10 | >10 | Indicator not available | |
| Poverty gap at $5.50 a day (2011 PPP) (% of population) | 40 | 11 | 41 | 6 | [70] |
| Poverty gap at national poverty lines (%) | 16 | 7 | 28 | 47 | [70] |
| Poverty headcount ratio at $1.90 a day (2011 PPP) (% of population) | 39 | 8 | 45 | 6 | [70] |
| Poverty headcount ratio at $3.20 a day (2011 PPP) (% of population) | 40 | 11 | 41 | 6 | [70] |
| Poverty headcount ratio at $5.50 a day (2011 PPP) (% of population) | 40 | 11 | 41 | 6 | [70] |
| Poverty headcount ratio at national poverty lines (% of population) | 27 | 8 | 40 | 23 | [70] |
| GDP per capita, PPP (constant 2011 international $) | 46 | 9 | 43 | 0 | [70] |
| GDP per capita, PPP (current international $) | 46 | 9 | 43 | 0 | [70] |
| HDI | 41 | 12 | 45 | 0 | [71] |
| GDP per capita, current prices (Purchasing power parity; international dollars per capita) | 61 | 6 | 31 | 0 | [72] |
| GDP per capita (1990 Int. GK$) | 97 | 0 | 0 | 1 | [73] |
| **Educational** | | | | | |
| Adjusted net enrollment rate, primary (% of primary school age children) | 25 | 12 | 56 | 5 | [70] |
| Educational attainment, at least completed upper secondary, population 25+, female (%) (cumulative) | 27 | 15 | 46 | 10 | [70] |
| Educational attainment, at least completed upper secondary, population 25+, male (%) (cumulative) | 27 | 15 | 46 | 10 | [70] |
| Educational attainment, at least completed upper secondary, population 25+, total (%) (cumulative) | 29 | 15 | 44 | 10 | [70] |
| **Employment** | | | | | [70] |
| Unemployment, total (% of total labor force) (modeled ILO estimate) | 45 | 9 | 44 | 0 | [70] |
| Unemployment, total (% of total labor force) (national estimate) | 63 | 18 | 17 | 0 | [70] |
| Employment to population ratio, 15+, female (%) (modeled ILO estimate) | 45 | 9 | 44 | 0 | [70] |

**Table S2** (continued)

| | Number of years difference between the year in which the indicator was available and the study year | | | | Data source |
|---|---|---|---|---|---|
| | <5 | 5-10 | >10 | Indicator not available | |
| Employment to population ratio, 15+, female (%) (national estimate) | 32 | 16 | 37 | 13 | [70] |
| Employment to population ratio, ages 15-24, female (%) (modeled ILO estimate) | 45 | 9 | 44 | 0 | [70] |
| Employment to population ratio, ages 15-24, female (%) (national estimate) | 18 | 10 | 52 | 18 | [70] |
| **Health-related** | | | | | |
| Low-birthweight babies (% of births) | 32 | 11 | 55 | 0 | [70] |
| Prevalence of undernourishment (% of population) | 39 | 6 | 32 | 21 | [70] |
| Prevalence of underweight, weight for age (% of children under 5) | 48 | 20 | 26 | 4 | [70] |
| Health expenditure, total (% of GDP) | 36 | 13 | 49 | 0 | [70] |
| Immunization, DPT (% of children ages 12-23 months) | 62 | 6 | 30 | 0 | [70] |
| Immunization, HepB3 (% of one-year-old children) | 28 | 10 | 53 | 7 | [70] |
| Immunization, measles (% of children ages 12-23 months) | 60 | 5 | 33 | 0 | [70] |
| Exclusive breastfeeding (% of children under 6 months) | 31 | 10 | 42 | 15 | [70] |
| Physicians (per 1,000 people) | 95 | 1 | 2 | 0 | [70] |
| Number of doctors' consultations | 13 | 3 | 9 | 73 | [74] |
| HiB vaccination coverage | 17 | 13 | 66 | 2 | [75] |
| **Housing** | | | | | |
| Average number of people per room in occupied housing unit | 5 | 1 | 6 | 86 | [76] |
| Population living in slums (% of urban population) | 39 | 3 | 33 | 23 | [70] |
| Number of households All households - Per capita | 12 | 4 | 45 | 37 | [77] |
| Number of households 1 person - Per capita | 9 | 2 | 45 | 42 | [77] |
| Number of households 1 person - Proportion over All households | 9 | 2 | 45 | 42 | [77] |
| Number of households 2 persons - Per capita | 9 | 2 | 45 | 42 | [77] |
| Number of households 2 persons - Proportion over All households | 9 | 2 | 45 | 42 | [77] |

**Table S2** (continued)

| | Number of years difference between the year in which the indicator was available and the study year | | | | Data source |
|---|---|---|---|---|---|
| | **<5** | **5-10** | **>10** | **Indicator not available** | |
| Number of households 3 persons - Per capita | 9 | 2 | 45 | 42 | [77] |
| Number of households 3 persons - Proportion over All households | 9 | 2 | 45 | 42 | [77] |
| Number of households 4 persons - Per capita | 9 | 2 | 45 | 42 | [77] |
| Number of households 4 persons - Proportion over All households | 9 | 2 | 45 | 42 | [77] |
| Number of households 5 persons - Per capita | 9 | 2 | 45 | 42 | [77] |
| Number of households 5 persons - Proportion over All households | 9 | 2 | 45 | 42 | [77] |
| Number of households 6 persons and over - Per capita | 9 | 2 | 42 | 45 | [77] |
| Number of households 6 persons and over - Proportion over All households | 9 | 2 | 42 | 45 | [77] |
| Total population in a household, both sexes | 2 | 2 | 22 | 72 | [78] |

**Table S3:** Number of missing indicators for each of the 98 studies

| Study Name | Number of missing indicator values |
|---|---|
| Benin, 1993[1] | 15 |
| Burkina Faso, 2007-8[6] | 2 |
| Congo, <1991[10] | 18 |
| Cote d'Ivoire, 1975[15] | 18 |
| Cote d'Ivoire, 1985-6[19] | 18 |
| Democratic Republic of the Congo (Kikwit), 2008-9[22] | 20 |
| Democratic Republic of the Congo (Mikalayi), 2008-9[22] | 20 |
| Democratic Republic of the Congo (Tshikapa), 2008-9[22] | 20 |
| Democratic Republic of the Congo (Vanga), 2008-9[22] | 20 |
| Ethiopia, 1981[38] | 3 |
| Ethiopia (Addis Ababa), 1994[42] | 3 |
| Gabon, 1985[45] | 20 |
| Ghana, 1997[49] | 2 |
| Kenya (Eldoret), 2005[54] | 18 |
| Kenya (Kilifi), 1996-9[56, 57] | 18 |
| Madagascar, 1990-1995[58] | 19 |
| Mozambique, 2002[59] | 16 |
| Nigeria, <1978[60] | 21 |
| Nigeria, <2002[61] | 21 |
| Nigeria, 2007-8[63] | 21 |
| Senegal, 1996-2001[65] | 16 |
| South Africa, 2003[66] | 1 |
| Tanzania (Mwanza), 2012-13[67] | 16 |
| Zambia, 1979-80[68] | 5 |
| Argentina (rural), 1967-68[2] | 4 |
| Argentina (urban), 1967-68[2] | 4 |
| Argentina (Mar de Plata), 1981[11] | 4 |
| Brazil, 1967-68[2] | 2 |
| Brazil, 1987[20] | 2 |
| Brazil (Parana), 1996-8[23] | 2 |
| Canada, <1967[4] | 6 |
| Chile (rural), 1967-68[2] | 3 |
| Chile (Santiago), 1967-68[2] | 3 |
| Haiti, 2002[39] | 17 |
| Jamaica (Kingston), 1967-68[2] | 6 |
| Jamaica (rural), 1967-68[2] | 6 |
| Mexico, 1987-88[50] | 1 |
| Mexico, 1989[55] | 1 |
| Panama (Panama City), 1967-68[2] | 17 |

| | |
|---|---|
| Panama (rural), 1967-68[2] | 17 |
| Peru (Lima), 1967-68[2] | 3 |
| Peru (rural), 1967-68[2] | 3 |
| Peru, 2003[62] | 3 |
| Trinidad, 1966-7[64] | 7 |
| Trinidad (rural), 1967-8[2] | 7 |
| Trinidad (Port au Spain), 1967-68[2] | 7 |
| Uruguay (rural) 1967-68[2] | 3 |
| Uruguay (urban) 1967-68[2] | 3 |
| USA (Atlanta), <1967[4] | 6 |
| USA (Houston), <1967[4] | 6 |
| Bahrain, 1981[3] | 30 |
| Iran, 1993-95[7] | 5 |
| Jordan, 1982-3[12] | 16 |
| Kuwait, <1978[16] | 30 |
| Lebanon, 1980-81[21] | 16 |
| Morocco, 1969-1970[24] | 19 |
| Pakistan, <1997[27] | 16 |
| Pakistan, 1999-2004[31] | 16 |
| Saudi Arabia, 1989[34] | 28 |
| Saudi Arabia, 1992-93[40] | 28 |
| Tunisia, <1970[43] | 17 |
| Yemen, 1985[46] | 19 |
| Yemen, 2002-03[51] | 19 |
| Czech Republic, <1967[4] | 5 |
| Denmark, <1967[4] | 21 |
| Denmark, 1983[13] | 21 |
| East Germany, 1990[17] | 5 |
| England, <1967[4] | 6 |
| England, 1986-87[25] | 6 |
| Finland, 1979[28] | 7 |
| France, <1967[4] | 20 |
| Kyrgyzstan, 2001[35] | 3 |
| Romania, <1989[41] | 4 |
| Turkey, 1998[44] | 14 |
| Turkey, 2003-04[47] | 14 |
| Turkey, 2005[52] | 14 |
| Bangladesh, 2004-05[5] | 4 |
| India (rural Delhi), 1968[8] | 3 |
| India (urban Delhi), 1968[8] | 3 |
| India (Chandrigarh), 1972-3[8] | 3 |
| India (Lucknow), 1972-3[8] | 3 |
| India (Calcutta), 1976[26] | 3 |
| India (Delhi), <1987[29] | 3 |

| | |
|---|---|
| India (Delhi), <1990[32] | 3 |
| India (rural Vellore), 1999-2000[36] | 3 |
| India (urban Vellore), 1999-2000[36] | 3 |
| Indonesia, 2007 (*S Reef, personal communication, March 2015*) | 16 |
| Nepal, 2008[48] | 4 |
| Thailand, 1978[53] | 5 |
| Australia, <1967[4] | 6 |
| Cambodia, 2012[9] | 25 |
| China, 1979-80[14] | 9 |
| Fiji, <1973[18] | 19 |
| Japan (Sapporo), <1967[4] | 7 |
| Japan (Ohtsu), <1967[4] | 7 |
| Malaysia, <1972[30] | 3 |
| Singapore, 1975-79[33] | 19 |
| Central Vietnam, 2009-2010[37] | 3 |

### S3 Details of 4-fold cross-validation – Dealing with missing indicators in simple linear regression and random forest analyses

Pseudocode description

The following piece of pseudocode describes the practical details of the 4-fold cross-validation experiment and the calculation of the MSE metrics of the $R_0$ predictions from the linear regression and the random forest prediction algorithms against the values estimated using seroprevalence data as well as the calculation of the corresponding MSE metrics for the case where $R_0$ is calculated using the default method against the values estimated using country-specific seroprevalence data.

```
1. RandomState = 0.  Initialize RandomState variable which is used
   as the seed for the pseudo-random split of the 98 studies to 4
   folds (point 3.a)
2. GoodStatesList = [empty list].  Initialize GoodStatesList in
   which we keep the seeds that result in splits to 4 folds that are
   suitable for running the experiment (see point 3.b.i below)
3. while length(GoodStatesList) ≤ 10
   a. Split the 98 studies in 4 folds (24 or 25 studies in each)
      using RandomState as the seed
   b. Check whether for each one of the 66 indicators there is at
      least one study in each fold with a valid (non-missing) value
      of the indicator:
      i.  If Yes: append RandomState to GoodStatesList and continue
          to the following step 3.c
     ii.  If No: then increase the value of RandomState by 1 and
          jump to step 3.a
   Simple linear regression component:
   c. for indicator, i, ranging across the 66 indicators
      i.  for testfold, t, ranging from 1 to 4
          A.  Use those studies in testfold that have a valid value
              for indicator i as the test set (denote their number
              with N_test) and those studies in the other 3 folds
              that have a valid value for indicator i as the train
              set (denote their number with N_train). As a result of
              check in step 3.b.i it is guaranteed that N_test ≥ 1
              and N_train ≥ 3. (A simple explanation of this is given
              in the toy-case example further below). The number of
              studies in the test and train set for each indicator
              in each fold and each different split to folds are
              given in the Table S4 further below
          B.  Fit linear regression line to the N_train studies in the
              training set using the value R_0 of the basic
              reproduction number as estimated using seroprevalence
              data
```

C. Use the fitted line calculated in step 3.c.i.B to predict the basic reproduction number $\hat{R}_0$ for each of the $N_{test}$ studies in the test set.

D. Compute the mean squared error for the studies in the test set, $MSE_{LR}(t,i)$ as

$$MSE_{LR}(t,i) = \sum_{j=1}^{N_{test}} \frac{\left(\hat{R}_{0_j} - R_{0_j}\right)^2}{N_{test}}$$

where $\hat{R}_{0_j}$ and $R_{0_j}$ denote the basic reproduction number for the j^th study of the test set predicted using linear regression line of step 3.c.i.B and estimated using seroprevalence data respectively.

E. Compute the mean squared error for the studies in the test set, $MSE_{LR,def}(t,i)$ as

$$MSE_{LR,def}(t,i) = \sum_{j=1}^{N_{test}} \frac{\left(\hat{R}_{0,def_j} - R_{0_j}\right)^2}{N_{test}}$$

where $\hat{R}_{0,def_j}$ and $R_{0_j}$ denote the basic reproduction number for the j^th study of the test set predicted using the default method and estimated using country-specific seroprevalence respectively

ii. Average the values of MSE over the 4 folds

A. Compute the mean of $MSE_{LR}(t, i)$ over the four *testfold* repetitions to get $MSE_{LR_{mean}}(i)$ (linear regression prediction):

$$MSE_{LR_{mean}}(i) = \sum_{t=1}^{4} MSE_{LR}(t,i)/4$$

B. Compute the mean of $MSE_{LR,def}(t, i)$ over the four *testfold* repetions to get $MSE_{LR,def_{mean}}(i)$ (regional average prediction):

$$MSE_{LR,def_{mean}}(i) = \sum_{t=1}^{4} \frac{MSE_{LR,def}(t,i)}{4}$$

***Random forest component:***

d. for *indicatorset*, $I_k$, $k = 1, 2, 3, 4, 5$ ranging across the 5 subsets of indicators described in the text (25 indicators that have no missing values, 43 indicators that have up to 10 missing values, etc.)

i. for *testfold*, $t$, ranging from 1 to 4

A. Use those studies in *testfold* that have valid values for all the indicators in *indicatorset*, $I_k$ as the test set (denote their number with $N_{test}$) and those studies in the other 3 folds that have valid values for all the indicators in *indicatorset*, $I_k$ as the training set (denote their number with $N_{train}$). Note that the check of step 3.b.i guarantees neither that $N_{test} > 0$ nor that $N_{train} > 0$. A simple explanation of this is given in the toy-case example further below. The number of studies in the test and train set for each indicators set $I_k$ in each fold and each different split to folds are given in Table S4 further below

B. If any of $N_{test}$ or $N_{train}$ is 0 raise a flag and stop the execution

C. Train a random forest with the $N_{train}$ studies of the train set

D. Use the trained random forest of step 3.d.i.C to predict the basic reproduction number $\hat{R}_0$ for each of the $N_{test}$ studies in the test set.

E. Compute the mean squared error for the studies in the test set $MSE_{RF}(t, I_k)$ as

$$MSE_{RF}(t, I_k) = \sum_{j=1}^{N_{test}} \frac{\left(\hat{R}_{0_j} - R_{0_j}\right)^2}{N_{test}}$$

where $\hat{R}_{0_j}$ and $R_{0_j}$ denote the basic reproduction number for the j[th] study of the test set predicted using the random forest of step 3.d.i.C and estimated using seroprevalence data respectively

F. Compute the mean squared error for the studies in the test set $MSE_{RF,def}(t, I_k)$ as

$$MSE_{RF,def}(t, I_k) = \sum_{j=1}^{N_{test}} \frac{\left(\hat{R}_{0,def_j} - R_{0_j}\right)^2}{N_{test}}$$

where $\hat{R}_{0,def_j}$ and $R_{0_j}$ is the basic reproduction number for the j[th] study of the test set predicted using the default method and estimated using country-specific seroprevalence data respectively

ii. Average the values of MSE over the 4 folds

A. Compute the mean of $MSE_{RF}(t, I_k)$ over the four *testfold* repetions to get $MSE_{RF_{mean}}(I_k)$ (random forest prediction):

$$MSE_{RF_{mean}}(I_k) = \sum_{t=1}^{4} MSE_{RF}(t, I_k)/4$$

B. Compute the mean of $MSE_{RF,def}(t, I_k)$ over the four *testfold* repetions to get $MSE_{RF,def_{mean}}(I_k)$ (regional average prediction):

$$MSE_{RF,def_{mean}}(I_k) = \sum_{t=1}^{4} \frac{MSE_{RF,def}(t, I_k)}{4}$$

e. Store values of $MSE_{LR_{mean}}(i)$ and $MSE_{LR,df_{mean}}(i)$for each of the 66 indicators and $MSE_{RF_{mean}}(I_k)$ and $MSE_{RF,def_{mean}}(I_k)$ for each indicator set $I_k$ (k = 1, .., 5)

f. Increase the value of *RandomState* by 1 and move to next iteration of while-loop in step 3

Missing values example

In the following we use a toy-case example to highlight the implications that missing indicator values have in the k-fold cross-validation experiments for the simple linear regression and the random forest prediction algorithms. We consider a simplistic scenario in which there are 9 data points (in our case studies) and 5 features (in our case socio-economic indicators). We take example case in which the missing values are distributed as follows:

- Indicator 1 has no missing values
- Indicators 2 and 3 are missing for Study 9
- Indicator 4 is missing in Studies 8 and 9
- Indicator 5 is missing in Studies 1, 5, 6, 7, and 8

We consider the case in which a 3-fold cross-validation is used. For a chosen split of the 9 studies to 3 folds, in order for cross-validation experiment to be feasible in the simple linear regression case, it is required that for every one of the indicators there exists at least one study in each one of the folds that has a valid value. Such an example of a split to folds where the simple linear regression cross-validation experiment is feasible for each one of the indicators is shown in Fig S1 (labelled as ‘Example 1’). Furthermore, in this particular split to folds it can also be seen that there exists a study in each fold which has valid values for all indicators. This practically means that the cross-validation experiment is also feasible for the random forest case in which one predictor is fitted to all the indicators together.

In a second example of split to folds, shown in Fig S1 (labelled as ‘Example 2’), it is also true that for every one of the indicators there is at least one study in each one of the folds which has a valid value. However, in the split of Example 2, it can be seen that Fold 3 does contain a study which has valid values for all indicators. Hence in this split the cross-validation experiment is feasible for the simple linear regression algorithm on each of the indicators but not for the random forest algorithm which is trained and tested on all the indicators together. The random forest cross-validation experiment is however feasible in both splits of Fig S1 if we restrict the random forest training/testing to a subset of indicators, e.g. to indicators 1 to 4. As it can be seen in Fig S1 if we restrict the random forest to train/testing using indicators 1 to 4, then in example 2 split 1 has studies 1, 2 and 4 with full values, and the same for studies 3 and 7 in split 2 and studies 5 and 6 in split 3..

**Fig S1:** Examples of two different splits to folds for the toy-case cross-validation scenario described in section S3. The ‘x’ marks denote missing values

Example 1

| Fold | Study | Indicator | | | | |
|---|---|---|---|---|---|---|
| | | 1 | 2 | 3 | 4 | 5 |
| 1 | 4 | | | | | |
| | 9 | | x | x | x | |
| | 1 | | | | | x |
| 2 | 3 | | | | | |
| | 8 | | | | x | x |
| | 6 | | | | | x |
| 3 | 5 | | | | | x |
| | 2 | | | | | |
| | 7 | | | | | x |

Example 2

| Fold | Study | Indicator | | | | |
|---|---|---|---|---|---|---|
| | | 1 | 2 | 3 | 4 | 5 |
| 1 | 4 | | | | | |
| | 2 | | | | | |
| | 1 | | | | | x |
| 2 | 7 | | | | | x |
| | 8 | | | | x | x |
| | 3 | | | | | |
| 3 | 5 | | | | | x |
| | 9 | | x | x | x | |
| | 6 | | | | | x |

Overall, as can be seen in Fig S1, in the example considered here there are only three studies that have valid values for all indicators so only a few splits to folds have exactly one of those 3 studies in each fold. By using a smaller set of indicators this condition is relaxed. For instance, there are 7 studies that have valid values for the subset containing indicators 1 to 4. Consequently, the cross-validation experiment is feasible for the random forest in more splits to folds.

As is shown in Table S3, in the actual indicator data that we use for the results presented in this work, there is no study having a valid value for all indicators. Hence, in order to make possible the random forest cross-validation experiment feasible we chose to run it on subsets of indicators as described in the main text. As can be seen in Table S4, for the 5 subsets of indicators described in the main text it was the case that in each of the 10 splits to folds chosen for the simple linear regression experiment there was at least one study in each fold which had valid values for all indicators in the chosen indicators subset. This can be seen in Table S4

**Table S4:** Number of studies with a valid indicator value in each fold for each of the 10 repetitions of the 4-fold cross validation experiment. The indicators order for the Simple Linear Regression case is the same as in **Table S2** and the order of indicators subsets for the Random Forest case is the same as in the main text. The row labelled 'Seed' gives the seed numbers used for each of the 10 split to folds (omitted seed numbers corresponds to splits that did not have at least one study with a valid (non-missing) value in each fold for each one of the 66 indicators.

| Split | 1 | | | | 2 | | | | 3 | | | | 4 | | | | 5 | | | | 6 | | | | 7 | | | | 8 | | | | 9 | | | | 10 | | | |
|---|---|---|---|---|---|---|---|---|---|---|---|---|---|---|---|---|---|---|---|---|---|---|---|---|---|---|---|---|---|---|---|---|---|---|---|---|---|---|---|---|
| Seed | 0 | | | | 8 | | | | 9 | | | | 10 | | | | 11 | | | | 12 | | | | 13 | | | | 14 | | | | 16 | | | | 17 | | | |
| Fold | 1 | 2 | 3 | 4 | 1 | 2 | 3 | 4 | 1 | 2 | 3 | 4 | 1 | 2 | 3 | 4 | 1 | 2 | 3 | 4 | 1 | 2 | 3 | 4 | 1 | 2 | 3 | 4 | 1 | 2 | 3 | 4 | 1 | 2 | 3 | 4 | 1 | 2 | 3 | 4 |
| Indicator | Simple Linear Regression | | | | | | | | | | | | | | | | | | | | | | | | | | | | | | | | | | | | | | | |
| 1 | 25 | 25 | 24 | 24 | 25 | 25 | 24 | 24 | 25 | 25 | 24 | 24 | 25 | 25 | 24 | 24 | 25 | 25 | 24 | 24 | 25 | 25 | 24 | 24 | 25 | 25 | 24 | 24 | 25 | 25 | 24 | 24 | 25 | 25 | 24 | 24 | 25 | 25 | 24 | 24 |
| 2 | 25 | 25 | 24 | 24 | 25 | 25 | 24 | 24 | 25 | 25 | 24 | 24 | 25 | 25 | 24 | 24 | 25 | 25 | 24 | 24 | 25 | 25 | 24 | 24 | 25 | 25 | 24 | 24 | 25 | 25 | 24 | 24 | 25 | 25 | 24 | 24 | 25 | 25 | 24 | 24 |
| 3 | 25 | 25 | 24 | 24 | 25 | 25 | 24 | 24 | 25 | 25 | 24 | 24 | 25 | 25 | 24 | 24 | 25 | 25 | 24 | 24 | 25 | 25 | 24 | 24 | 25 | 25 | 24 | 24 | 25 | 25 | 24 | 24 | 25 | 25 | 24 | 24 | 25 | 25 | 24 | 24 |
| 4 | 25 | 25 | 24 | 24 | 25 | 25 | 24 | 24 | 25 | 25 | 24 | 24 | 25 | 25 | 24 | 24 | 25 | 25 | 24 | 24 | 25 | 25 | 24 | 24 | 25 | 25 | 24 | 24 | 25 | 25 | 24 | 24 | 25 | 25 | 24 | 24 | 25 | 25 | 24 | 24 |
| 5 | 25 | 25 | 24 | 24 | 25 | 25 | 24 | 24 | 25 | 25 | 24 | 24 | 25 | 25 | 24 | 24 | 25 | 25 | 24 | 24 | 25 | 25 | 24 | 24 | 25 | 25 | 24 | 24 | 25 | 25 | 24 | 24 | 25 | 25 | 24 | 24 | 25 | 25 | 24 | 24 |
| 6 | 25 | 25 | 24 | 24 | 25 | 25 | 24 | 24 | 25 | 25 | 24 | 24 | 25 | 25 | 24 | 24 | 25 | 25 | 24 | 24 | 25 | 25 | 24 | 24 | 25 | 25 | 24 | 24 | 25 | 25 | 24 | 24 | 25 | 25 | 24 | 24 | 25 | 25 | 24 | 24 |
| 7 | 25 | 25 | 24 | 24 | 25 | 25 | 24 | 24 | 25 | 25 | 24 | 24 | 25 | 25 | 24 | 24 | 25 | 25 | 24 | 24 | 25 | 25 | 24 | 24 | 25 | 25 | 24 | 24 | 25 | 25 | 24 | 24 | 25 | 25 | 24 | 24 | 25 | 25 | 24 | 24 |
| 8 | 25 | 25 | 24 | 24 | 25 | 25 | 24 | 24 | 25 | 25 | 24 | 24 | 25 | 25 | 24 | 24 | 25 | 25 | 24 | 24 | 25 | 25 | 24 | 24 | 25 | 25 | 24 | 24 | 25 | 25 | 24 | 24 | 25 | 25 | 24 | 24 | 25 | 25 | 24 | 24 |
| 9 | 25 | 25 | 24 | 24 | 25 | 25 | 24 | 24 | 25 | 25 | 24 | 24 | 25 | 25 | 24 | 24 | 25 | 25 | 24 | 24 | 25 | 25 | 24 | 24 | 25 | 25 | 24 | 24 | 25 | 25 | 24 | 24 | 25 | 25 | 24 | 24 | 25 | 25 | 24 | 24 |
| 10 | 25 | 25 | 24 | 24 | 25 | 25 | 24 | 24 | 25 | 25 | 24 | 24 | 25 | 25 | 24 | 24 | 25 | 25 | 24 | 24 | 25 | 25 | 24 | 24 | 25 | 25 | 24 | 24 | 25 | 25 | 24 | 24 | 25 | 25 | 24 | 24 | 25 | 25 | 24 | 24 |
| 11 | 25 | 25 | 24 | 24 | 25 | 25 | 24 | 24 | 25 | 25 | 24 | 24 | 25 | 25 | 24 | 24 | 25 | 25 | 24 | 24 | 25 | 25 | 24 | 24 | 25 | 25 | 24 | 24 | 25 | 25 | 24 | 24 | 25 | 25 | 24 | 24 | 25 | 25 | 24 | 24 |
| 12 | 25 | 25 | 24 | 24 | 25 | 25 | 24 | 24 | 25 | 25 | 24 | 24 | 25 | 25 | 24 | 24 | 25 | 25 | 24 | 24 | 25 | 25 | 24 | 24 | 25 | 25 | 24 | 24 | 25 | 25 | 24 | 24 | 25 | 25 | 24 | 24 | 25 | 25 | 24 | 24 |
| 13 | 23 | 23 | 23 | 23 | 24 | 22 | 24 | 22 | 23 | 23 | 22 | 24 | 22 | 24 | 22 | 24 | 23 | 24 | 22 | 23 | 24 | 24 | 23 | 21 | 24 | 25 | 24 | 19 | 25 | 24 | 22 | 21 | 22 | 25 | 22 | 23 | 24 | 23 | 23 | 22 |
| 14 | 23 | 23 | 23 | 23 | 24 | 22 | 24 | 22 | 23 | 23 | 22 | 24 | 22 | 24 | 22 | 24 | 23 | 24 | 22 | 23 | 24 | 24 | 23 | 21 | 24 | 25 | 24 | 19 | 25 | 24 | 22 | 21 | 22 | 25 | 22 | 23 | 24 | 23 | 23 | 22 |
| 15 | 23 | 23 | 23 | 23 | 24 | 22 | 24 | 22 | 23 | 23 | 22 | 24 | 22 | 24 | 22 | 24 | 23 | 24 | 22 | 23 | 24 | 24 | 23 | 21 | 24 | 25 | 24 | 19 | 25 | 24 | 22 | 21 | 22 | 25 | 22 | 23 | 24 | 23 | 23 | 22 |
| 16 | 23 | 23 | 23 | 23 | 24 | 22 | 24 | 22 | 23 | 23 | 22 | 24 | 22 | 24 | 22 | 24 | 23 | 24 | 22 | 23 | 24 | 24 | 23 | 21 | 24 | 25 | 24 | 19 | 25 | 24 | 22 | 21 | 22 | 25 | 22 | 23 | 24 | 23 | 23 | 22 |
| 17 | 23 | 23 | 23 | 23 | 24 | 22 | 24 | 22 | 23 | 23 | 22 | 24 | 22 | 24 | 22 | 24 | 23 | 24 | 22 | 23 | 24 | 24 | 23 | 21 | 24 | 25 | 24 | 19 | 25 | 24 | 22 | 21 | 22 | 25 | 22 | 23 | 24 | 23 | 23 | 22 |
| 18 | 23 | 23 | 23 | 23 | 24 | 22 | 24 | 22 | 23 | 23 | 22 | 24 | 22 | 24 | 22 | 24 | 23 | 24 | 22 | 23 | 24 | 24 | 23 | 21 | 24 | 25 | 24 | 19 | 25 | 24 | 22 | 21 | 22 | 25 | 22 | 23 | 24 | 23 | 23 | 22 |
| 19 | 23 | 23 | 23 | 23 | 24 | 22 | 24 | 22 | 23 | 23 | 22 | 24 | 22 | 24 | 22 | 24 | 23 | 24 | 22 | 23 | 24 | 24 | 23 | 21 | 24 | 25 | 24 | 19 | 25 | 24 | 22 | 21 | 22 | 25 | 22 | 23 | 24 | 23 | 23 | 22 |
| 20 | 15 | 13 | 10 | 13 | 11 | 13 | 12 | 15 | 14 | 12 | 9 | 16 | 11 | 15 | 11 | 14 | 14 | 14 | 11 | 12 | 16 | 6 | 15 | 14 | 12 | 15 | 13 | 11 | 12 | 14 | 13 | 12 | 13 | 14 | 10 | 14 | 12 | 14 | 12 | 13 |

| | | | | | | | | | | | | | | | | | | | | | | | | | | | | | | | | | | | | | | | | |
|---|---|---|---|---|---|---|---|---|---|---|---|---|---|---|---|---|---|---|---|---|---|---|---|---|---|---|---|---|---|---|---|---|---|---|---|---|---|---|---|---|
| **21** | 23 | 23 | 23 | 23 | 24 | 22 | 24 | 22 | 23 | 23 | 22 | 24 | 22 | 24 | 22 | 24 | 23 | 24 | 22 | 23 | 24 | 24 | 23 | 21 | 24 | 25 | 24 | 19 | 25 | 24 | 22 | 21 | 22 | 25 | 22 | 23 | 24 | 23 | 23 | 22 |
| **22** | 23 | 23 | 23 | 23 | 24 | 22 | 24 | 22 | 23 | 23 | 22 | 24 | 22 | 24 | 22 | 24 | 23 | 24 | 22 | 23 | 24 | 24 | 23 | 21 | 24 | 25 | 24 | 19 | 25 | 24 | 22 | 21 | 22 | 25 | 22 | 23 | 24 | 23 | 23 | 22 |
| **23** | 23 | 23 | 23 | 23 | 24 | 22 | 24 | 22 | 23 | 23 | 22 | 24 | 22 | 24 | 22 | 24 | 23 | 24 | 22 | 23 | 24 | 24 | 23 | 21 | 24 | 25 | 24 | 19 | 25 | 24 | 22 | 21 | 22 | 25 | 22 | 23 | 24 | 23 | 23 | 22 |
| **24** | 19 | 18 | 20 | 18 | 18 | 20 | 17 | 20 | 22 | 19 | 14 | 20 | 16 | 22 | 18 | 19 | 21 | 17 | 20 | 17 | 20 | 17 | 19 | 19 | 20 | 20 | 21 | 14 | 18 | 20 | 19 | 18 | 19 | 20 | 16 | 20 | 16 | 21 | 19 | 19 |
| **25** | 25 | 25 | 24 | 24 | 25 | 25 | 24 | 24 | 25 | 25 | 24 | 24 | 25 | 25 | 24 | 24 | 25 | 25 | 24 | 24 | 25 | 25 | 24 | 24 | 25 | 25 | 24 | 24 | 25 | 25 | 24 | 24 | 25 | 25 | 24 | 24 | 25 | 25 | 24 | 24 |
| **26** | 25 | 25 | 24 | 24 | 25 | 25 | 24 | 24 | 25 | 25 | 24 | 24 | 25 | 25 | 24 | 24 | 25 | 25 | 24 | 24 | 25 | 25 | 24 | 24 | 25 | 25 | 24 | 24 | 25 | 25 | 24 | 24 | 25 | 25 | 24 | 24 | 25 | 25 | 24 | 24 |
| **27** | 25 | 25 | 24 | 24 | 25 | 25 | 24 | 24 | 25 | 25 | 24 | 24 | 25 | 25 | 24 | 24 | 25 | 25 | 24 | 24 | 25 | 25 | 24 | 24 | 25 | 25 | 24 | 24 | 25 | 25 | 24 | 24 | 25 | 25 | 24 | 24 | 25 | 25 | 24 | 24 |
| **28** | 25 | 25 | 24 | 24 | 25 | 25 | 24 | 24 | 25 | 25 | 24 | 24 | 25 | 25 | 24 | 24 | 25 | 25 | 24 | 24 | 25 | 25 | 24 | 24 | 25 | 25 | 24 | 24 | 25 | 25 | 24 | 24 | 25 | 25 | 24 | 24 | 25 | 25 | 24 | 24 |
| **29** | 24 | 25 | 24 | 24 | 25 | 25 | 24 | 23 | 25 | 24 | 24 | 24 | 25 | 25 | 23 | 24 | 25 | 25 | 23 | 24 | 25 | 25 | 24 | 23 | 25 | 25 | 23 | 24 | 25 | 24 | 24 | 24 | 25 | 25 | 24 | 23 | 25 | 25 | 24 | 23 |
| **30** | 24 | 25 | 22 | 22 | 24 | 21 | 24 | 24 | 24 | 24 | 22 | 23 | 24 | 24 | 23 | 22 | 23 | 25 | 24 | 21 | 25 | 24 | 22 | 22 | 25 | 24 | 22 | 22 | 23 | 25 | 22 | 23 | 25 | 24 | 22 | 22 | 23 | 22 | 24 | 24 |
| **31** | 24 | 23 | 22 | 19 | 21 | 21 | 23 | 23 | 21 | 23 | 22 | 22 | 23 | 23 | 23 | 19 | 23 | 23 | 22 | 20 | 21 | 24 | 23 | 20 | 19 | 21 | 24 | 24 | 22 | 22 | 21 | 23 | 23 | 21 | 22 | 22 | 21 | 22 | 23 | 22 |
| **32** | 24 | 23 | 22 | 19 | 21 | 21 | 23 | 23 | 21 | 23 | 22 | 22 | 23 | 23 | 23 | 19 | 23 | 23 | 22 | 20 | 21 | 24 | 23 | 20 | 19 | 21 | 24 | 24 | 22 | 22 | 21 | 23 | 23 | 21 | 22 | 22 | 21 | 22 | 23 | 22 |
| **33** | 24 | 23 | 22 | 19 | 21 | 21 | 23 | 23 | 21 | 23 | 22 | 22 | 23 | 23 | 23 | 19 | 23 | 23 | 22 | 20 | 21 | 24 | 23 | 20 | 19 | 21 | 24 | 24 | 22 | 22 | 21 | 23 | 23 | 21 | 22 | 22 | 21 | 22 | 23 | 22 |
| **34** | 25 | 25 | 24 | 24 | 25 | 25 | 24 | 24 | 25 | 25 | 24 | 24 | 25 | 25 | 24 | 24 | 25 | 25 | 24 | 24 | 25 | 25 | 24 | 24 | 25 | 25 | 24 | 24 | 25 | 25 | 24 | 24 | 25 | 25 | 24 | 24 | 25 | 25 | 24 | 24 |
| **35** | 25 | 25 | 24 | 24 | 25 | 25 | 24 | 24 | 25 | 25 | 24 | 24 | 25 | 25 | 24 | 24 | 25 | 25 | 24 | 24 | 25 | 25 | 24 | 24 | 25 | 25 | 24 | 24 | 25 | 25 | 24 | 24 | 25 | 25 | 24 | 24 | 25 | 25 | 24 | 24 |
| **36** | 25 | 25 | 24 | 24 | 25 | 25 | 24 | 24 | 25 | 25 | 24 | 24 | 25 | 25 | 24 | 24 | 25 | 25 | 24 | 24 | 25 | 25 | 24 | 24 | 25 | 25 | 24 | 24 | 25 | 25 | 24 | 24 | 25 | 25 | 24 | 24 | 25 | 25 | 24 | 24 |
| **37** | 24 | 24 | 19 | 18 | 18 | 22 | 23 | 22 | 18 | 24 | 21 | 22 | 20 | 24 | 21 | 20 | 21 | 23 | 23 | 18 | 22 | 22 | 21 | 20 | 23 | 20 | 20 | 22 | 24 | 21 | 22 | 18 | 20 | 21 | 24 | 20 | 21 | 23 | 23 | 18 |
| **38** | 25 | 25 | 24 | 24 | 25 | 25 | 24 | 24 | 25 | 25 | 24 | 24 | 25 | 25 | 24 | 24 | 25 | 25 | 24 | 24 | 25 | 25 | 24 | 24 | 25 | 25 | 24 | 24 | 25 | 25 | 24 | 24 | 25 | 25 | 24 | 24 | 25 | 25 | 24 | 24 |
| **39** | 21 | 24 | 19 | 16 | 16 | 22 | 23 | 19 | 17 | 21 | 21 | 21 | 20 | 22 | 19 | 19 | 19 | 23 | 21 | 17 | 20 | 22 | 20 | 18 | 23 | 19 | 19 | 19 | 24 | 20 | 20 | 16 | 19 | 20 | 23 | 18 | 21 | 21 | 22 | 16 |
| **40** | 25 | 25 | 24 | 24 | 25 | 25 | 24 | 24 | 25 | 25 | 24 | 24 | 25 | 25 | 24 | 24 | 25 | 25 | 24 | 24 | 25 | 25 | 24 | 24 | 25 | 25 | 24 | 24 | 25 | 25 | 24 | 24 | 25 | 25 | 24 | 24 | 25 | 25 | 24 | 24 |
| **41** | 20 | 21 | 18 | 18 | 19 | 20 | 19 | 19 | 20 | 20 | 18 | 19 | 20 | 22 | 16 | 19 | 18 | 21 | 21 | 17 | 21 | 19 | 19 | 18 | 22 | 19 | 19 | 17 | 19 | 19 | 20 | 19 | 21 | 21 | 16 | 19 | 19 | 19 | 19 | 20 |
| **42** | 22 | 25 | 24 | 23 | 24 | 25 | 22 | 23 | 24 | 24 | 23 | 23 | 22 | 25 | 24 | 23 | 25 | 23 | 23 | 23 | 24 | 25 | 22 | 23 | 25 | 24 | 23 | 22 | 23 | 25 | 23 | 23 | 24 | 24 | 24 | 22 | 24 | 24 | 23 | 23 |
| **43** | 25 | 25 | 24 | 24 | 25 | 25 | 24 | 24 | 25 | 25 | 24 | 24 | 25 | 25 | 24 | 24 | 25 | 25 | 24 | 24 | 25 | 25 | 24 | 24 | 25 | 25 | 24 | 24 | 25 | 25 | 24 | 24 | 25 | 25 | 24 | 24 | 25 | 25 | 24 | 24 |
| **44** | 25 | 25 | 24 | 24 | 25 | 25 | 24 | 24 | 25 | 25 | 24 | 24 | 25 | 25 | 24 | 24 | 25 | 25 | 24 | 24 | 25 | 25 | 24 | 24 | 25 | 25 | 24 | 24 | 25 | 25 | 24 | 24 | 25 | 25 | 24 | 24 | 25 | 25 | 24 | 24 |
| **45** | 21 | 23 | 23 | 24 | 22 | 24 | 22 | 23 | 24 | 24 | 21 | 22 | 22 | 25 | 21 | 23 | 23 | 22 | 23 | 23 | 24 | 22 | 22 | 23 | 25 | 22 | 24 | 20 | 23 | 23 | 23 | 22 | 23 | 23 | 22 | 23 | 24 | 24 | 20 | 23 |
| **46** | 25 | 25 | 24 | 24 | 25 | 25 | 24 | 24 | 25 | 25 | 24 | 24 | 25 | 25 | 24 | 24 | 25 | 25 | 24 | 24 | 25 | 25 | 24 | 24 | 25 | 25 | 24 | 24 | 25 | 25 | 24 | 24 | 25 | 25 | 24 | 24 | 25 | 25 | 24 | 24 |
| **47** | 20 | 22 | 21 | 20 | 22 | 20 | 19 | 22 | 23 | 22 | 18 | 20 | 21 | 22 | 20 | 20 | 22 | 21 | 22 | 18 | 22 | 22 | 19 | 20 | 23 | 21 | 21 | 18 | 19 | 22 | 20 | 22 | 22 | 22 | 18 | 21 | 20 | 21 | 20 | 22 |
| **48** | 25 | 25 | 24 | 24 | 25 | 25 | 24 | 24 | 25 | 25 | 24 | 24 | 25 | 25 | 24 | 24 | 25 | 25 | 24 | 24 | 25 | 25 | 24 | 24 | 25 | 25 | 24 | 24 | 25 | 25 | 24 | 24 | 25 | 25 | 24 | 24 | 25 | 25 | 24 | 24 |
| **49** | 8 | 7 | 6 | 4 | 5 | 8 | 8 | 4 | 6 | 4 | 9 | 6 | 5 | 6 | 8 | 6 | 4 | 6 | 8 | 7 | 4 | 8 | 8 | 5 | 5 | 6 | 8 | 6 | 9 | 6 | 4 | 6 | 5 | 6 | 7 | 7 | 6 | 4 | 12 | 3 |

| | | | | | | | | | | | | | | | | | | | | | | | | | | | | | | | | | | | | | | | | |
|---|---|---|---|---|---|---|---|---|---|---|---|---|---|---|---|---|---|---|---|---|---|---|---|---|---|---|---|---|---|---|---|---|---|---|---|---|---|---|---|---|
| **50** | 25 | 25 | 22 | 24 | 24 | 25 | 24 | 23 | 24 | 25 | 23 | 24 | 25 | 25 | 24 | 22 | 25 | 25 | 23 | 23 | 25 | 24 | 24 | 23 | 24 | 25 | 23 | 24 | 25 | 25 | 23 | 23 | 25 | 24 | 24 | 23 | 24 | 24 | 24 | 24 |
| **51** | 4 | 3 | 2 | 3 | 3 | 2 | 5 | 2 | 3 | 3 | 4 | 2 | 4 | 2 | 4 | 2 | 3 | 2 | 5 | 2 | 3 | 4 | 2 | 3 | 4 | 2 | 3 | 3 | 4 | 2 | 3 | 3 | 4 | 2 | 4 | 2 | 2 | 5 | 3 | 2 |
| **52** | 18 | 20 | 19 | 18 | 21 | 19 | 17 | 18 | 21 | 18 | 17 | 19 | 21 | 20 | 16 | 18 | 19 | 19 | 20 | 17 | 19 | 19 | 19 | 18 | 21 | 20 | 18 | 16 | 18 | 19 | 19 | 19 | 20 | 21 | 14 | 20 | 19 | 17 | 18 | 21 |
| **53** | 14 | 19 | 17 | 11 | 12 | 13 | 19 | 17 | 15 | 14 | 16 | 16 | 14 | 16 | 15 | 16 | 13 | 18 | 16 | 14 | 16 | 18 | 15 | 12 | 18 | 14 | 16 | 13 | 16 | 15 | 13 | 17 | 12 | 17 | 19 | 13 | 13 | 16 | 20 | 12 |
| **54** | 13 | 17 | 15 | 11 | 10 | 11 | 18 | 17 | 13 | 13 | 15 | 15 | 13 | 14 | 13 | 16 | 13 | 18 | 12 | 13 | 15 | 16 | 15 | 10 | 16 | 14 | 15 | 11 | 14 | 15 | 12 | 15 | 11 | 17 | 16 | 12 | 11 | 15 | 20 | 10 |
| **55** | 13 | 17 | 15 | 11 | 10 | 11 | 18 | 17 | 13 | 13 | 15 | 15 | 13 | 14 | 13 | 16 | 13 | 18 | 12 | 13 | 15 | 16 | 15 | 10 | 16 | 14 | 15 | 11 | 14 | 15 | 12 | 15 | 11 | 17 | 16 | 12 | 11 | 15 | 20 | 10 |
| **56** | 13 | 17 | 15 | 11 | 10 | 11 | 18 | 17 | 13 | 13 | 15 | 15 | 13 | 14 | 13 | 16 | 13 | 18 | 12 | 13 | 15 | 16 | 15 | 10 | 16 | 14 | 15 | 11 | 14 | 15 | 12 | 15 | 11 | 17 | 16 | 12 | 11 | 15 | 20 | 10 |
| **57** | 13 | 17 | 15 | 11 | 10 | 11 | 18 | 17 | 13 | 13 | 15 | 15 | 13 | 14 | 13 | 16 | 13 | 18 | 12 | 13 | 15 | 16 | 15 | 10 | 16 | 14 | 15 | 11 | 14 | 15 | 12 | 15 | 11 | 17 | 16 | 12 | 11 | 15 | 20 | 10 |
| **58** | 13 | 17 | 15 | 11 | 10 | 11 | 18 | 17 | 13 | 13 | 15 | 15 | 13 | 14 | 13 | 16 | 13 | 18 | 12 | 13 | 15 | 16 | 15 | 10 | 16 | 14 | 15 | 11 | 14 | 15 | 12 | 15 | 11 | 17 | 16 | 12 | 11 | 15 | 20 | 10 |
| **59** | 13 | 17 | 15 | 11 | 10 | 11 | 18 | 17 | 13 | 13 | 15 | 15 | 13 | 14 | 13 | 16 | 13 | 18 | 12 | 13 | 15 | 16 | 15 | 10 | 16 | 14 | 15 | 11 | 14 | 15 | 12 | 15 | 11 | 17 | 16 | 12 | 11 | 15 | 20 | 10 |
| **60** | 13 | 17 | 15 | 11 | 10 | 11 | 18 | 17 | 13 | 13 | 15 | 15 | 13 | 14 | 13 | 16 | 13 | 18 | 12 | 13 | 15 | 16 | 15 | 10 | 16 | 14 | 15 | 11 | 14 | 15 | 12 | 15 | 11 | 17 | 16 | 12 | 11 | 15 | 20 | 10 |
| **61** | 13 | 17 | 15 | 11 | 10 | 11 | 18 | 17 | 13 | 13 | 15 | 15 | 13 | 14 | 13 | 16 | 13 | 18 | 12 | 13 | 15 | 16 | 15 | 10 | 16 | 14 | 15 | 11 | 14 | 15 | 12 | 15 | 11 | 17 | 16 | 12 | 11 | 15 | 20 | 10 |
| **62** | 13 | 17 | 15 | 11 | 10 | 11 | 18 | 17 | 13 | 13 | 15 | 15 | 13 | 14 | 13 | 16 | 13 | 18 | 12 | 13 | 15 | 16 | 15 | 10 | 16 | 14 | 15 | 11 | 14 | 15 | 12 | 15 | 11 | 17 | 16 | 12 | 11 | 15 | 20 | 10 |
| **63** | 13 | 17 | 15 | 11 | 10 | 11 | 18 | 17 | 13 | 13 | 15 | 15 | 13 | 14 | 13 | 16 | 13 | 18 | 12 | 13 | 15 | 16 | 15 | 10 | 16 | 14 | 15 | 11 | 14 | 15 | 12 | 15 | 11 | 17 | 16 | 12 | 11 | 15 | 20 | 10 |
| **64** | 13 | 15 | 15 | 10 | 9 | 11 | 16 | 17 | 12 | 13 | 13 | 15 | 11 | 14 | 12 | 16 | 13 | 16 | 11 | 13 | 14 | 14 | 15 | 10 | 15 | 13 | 15 | 10 | 14 | 14 | 11 | 14 | 11 | 14 | 16 | 12 | 10 | 15 | 20 | 8 |
| **65** | 13 | 15 | 15 | 10 | 9 | 11 | 16 | 17 | 12 | 13 | 13 | 15 | 11 | 14 | 12 | 16 | 13 | 16 | 11 | 13 | 14 | 14 | 15 | 10 | 15 | 13 | 15 | 10 | 14 | 14 | 11 | 14 | 11 | 14 | 16 | 12 | 10 | 15 | 20 | 8 |
| **66** | 6 | 7 | 9 | 4 | 4 | 8 | 7 | 7 | 6 | 4 | 9 | 7 | 2 | 7 | 8 | 9 | 4 | 10 | 4 | 8 | 7 | 8 | 6 | 5 | 8 | 7 | 8 | 3 | 6 | 9 | 4 | 7 | 5 | 6 | 9 | 6 | 5 | 6 | 12 | 3 |
| **Indicator Subset** | **Random Forest** | | | | | | | | | | | | | | | | | | | | | | | | | | | | | | | | | | | | | | | |
| **1** | 25 | 25 | 24 | 24 | 25 | 25 | 24 | 24 | 25 | 25 | 24 | 24 | 25 | 25 | 24 | 24 | 25 | 25 | 24 | 24 | 25 | 25 | 24 | 24 | 25 | 25 | 24 | 24 | 25 | 25 | 24 | 24 | 25 | 25 | 24 | 24 | 25 | 25 | 24 | 24 |
| **2** | 16 | 19 | 18 | 16 | 16 | 15 | 20 | 18 | 17 | 18 | 15 | 19 | 17 | 22 | 16 | 14 | 18 | 19 | 17 | 15 | 19 | 19 | 18 | 13 | 17 | 18 | 20 | 14 | 18 | 18 | 16 | 17 | 18 | 17 | 17 | 17 | 17 | 16 | 18 | 18 |
| **3** | 13 | 17 | 13 | 9 | 10 | 11 | 17 | 14 | 12 | 14 | 11 | 15 | 14 | 16 | 13 | 9 | 13 | 16 | 15 | 8 | 14 | 17 | 14 | 7 | 15 | 12 | 16 | 9 | 15 | 15 | 11 | 11 | 12 | 14 | 13 | 13 | 12 | 13 | 16 | 11 |
| **4** | 9 | 12 | 8 | 3 | 4 | 5 | 12 | 11 | 8 | 8 | 6 | 10 | 9 | 8 | 9 | 6 | 8 | 9 | 10 | 5 | 9 | 8 | 10 | 5 | 9 | 7 | 11 | 5 | 10 | 9 | 4 | 9 | 8 | 8 | 7 | 9 | 5 | 7 | 14 | 6 |
| **5** | 6 | 9 | 3 | 2 | 1 | 1 | 8 | 10 | 5 | 4 | 4 | 7 | 6 | 4 | 5 | 5 | 6 | 8 | 4 | 2 | 8 | 2 | 7 | 3 | 5 | 5 | 7 | 3 | 5 | 6 | 3 | 6 | 5 | 6 | 3 | 6 | 3 | 5 | 8 | 4 |

### S4 Results – analyses of $R_0$ using the default approach and for pessimistic assumptions about contact

**Table S5:** Estimates of the mean square error associated with $R_0$ estimates. Column 2 shows the mean square error of $R_0$, as calculated using the default approach against $R_0$ calculated using country-specific seroprevalence data. Columns 2-7 hold the minimum, median, maximum 2.5th and 95.5th percentiles MSE values calculated using the 1000 bootstrap-derived values for $R_0$ for the force of infection from the region and the 1000 bootstrap-derived values calculated from country-specific seroprevalence data. The final column holds the MSE associated with $R_0$ from country-specific seroprevalence data vs $R_0$ from the regional point estimate of the force of infection

| | **$R_0$ from country-specific seroprevalence data vs default $R_0$ estimate** | **Min** | **2.5 %** | **Median** | **97.5%** | **Max** | **$R_0$ from country-specific seroprevalence data vs $R_0$ from regional point estimate of the force of infection** |
|---|---|---|---|---|---|---|---|
| **All countries** | 6.97 | 4.76 | 7.31 | 16.03 | 125.75 | 1513.39 | 8.31 |
| **Africa** | 1.03 | 0.73 | 1.01 | 2.16 | 16.23 | 65.14 | 2.23 |
| **Americas** | 8.72 | 4.62 | 6.60 | 12.04 | 298.38 | 1992.15 | 9.36 |
| **Eastern Mediterranean** | 5.01 | 2.73 | 3.88 | 6.46 | 75.69 | 3183.24 | 4.61 |
| **Europe** | 22.56 | 6.83 | 13.16 | 44.09 | 694.90 | 11359.21 | 28.10 |
| **South East Asia** | 4.71 | 1.33 | 2.38 | 7.65 | 133.70 | 440.50 | 6.14 |
| **Western Pacific** | 1.36 | 0.86 | 1.04 | 2.44 | 18.57 | 281.34 | 1.39 |

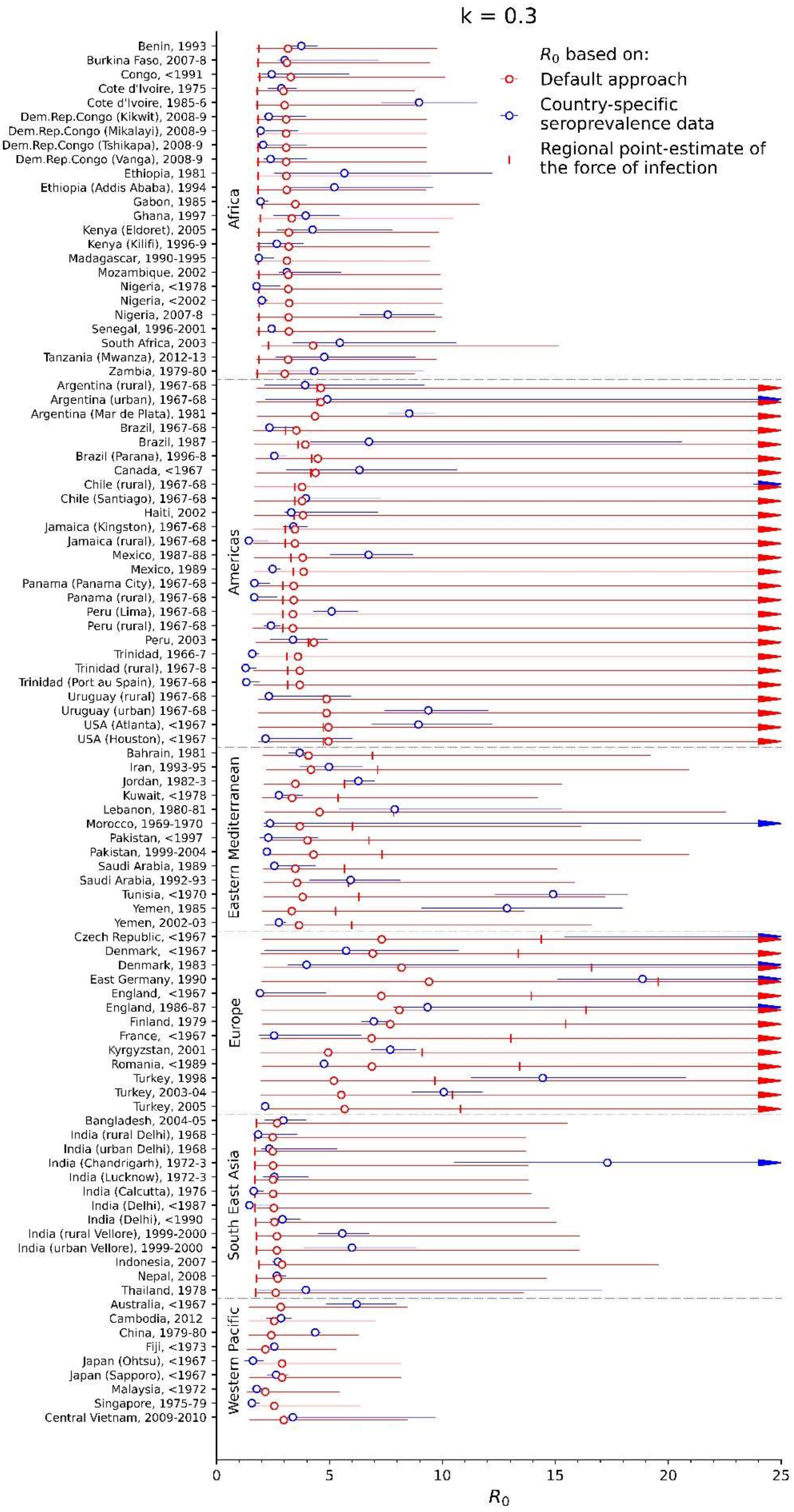

**Fig S2:** *Estimates of the basic reproduction number for rubella, as calculated using seroprevalence data (blue circles) and the default approach (red circles) for each study for pessimistic assumptions about the amount of contact between children and adults k=0.3). The vertical marks indicate $R_0$ as calculated using the regional point estimate for the force of infection.*

### S5 Results – Correlation between $R_0$ and the indicators

**Table S6:** Summary of the correlation coefficients and MIC for the association between the basic reproduction number and the indicators. ranked in decreasing order. The columns labelled "CC" hold the coefficient, with the 95% range obtained by bootstrapping; the column labelled "N" holds the number of data points used to calculate the coefficient.

| Rank | Pearson | | | Spearman | | | MIC | | |
|---|---|---|---|---|---|---|---|---|---|
| | **Indicator** | **CC** | **N** | **Indicator** | **CC** | **N** | **Indicator** | **CC** | **N** |
| 1 | Educational attainment, at least completed upper secondary, population 25+, total (%) (cumulative) | 0.4 (0.24, 0.48) | 88 | Number of households 5 persons - Proportion over All households | -0.45 (-0.5, -0.36) | 56 | Poverty gap at $1.90 a day (2011 PPP) (%) | 0.37 (0.21, 0.34) | 92 |
| 2 | Educational attainment, at least completed upper secondary, population 25+, female (%) (cumulative) | 0.4 (0.25, 0.47) | 88 | Number of households 5 persons - Per capita | -0.41 (-0.46, -0.32) | 56 | Poverty headcount ratio at $5.50 a day (2011 PPP) (% of population) | 0.36 (0.22, 0.37) | 92 |
| 3 | Educational attainment, at least completed upper secondary, population 25+, male (%) (cumulative) | 0.39 (0.22, 0.47) | 88 | Number of households 6 persons and over - Per capita | -0.33 (-0.41, -0.22) | 53 | Crude death rate per 1000 population | 0.35 (0.2, 0.37) | 98 |
| 4 | Number of households 5 persons - Proportion over All households | -0.34 (-0.42, -0.19) | 56 | Physicians (per 1,000 people) | 0.32 (0.22, 0.37) | 98 | Life expectancy at birth (both sexes) | 0.34 (0.2, 0.33) | 98 |
| 5 | Physicians (per 1,000 people) | 0.33 (0.16, 0.41) | 98 | Prevalence of underweight, weight for age (% of children under 5) | -0.31 (-0.34, -0.21) | 94 | Poverty gap at $3.20 a day (2011 PPP) (% of population) | 0.34 (0.22, 0.36) | 92 |
| 6 | Proportion of the population aged 65+ | 0.32 (0.2, 0.51) | 98 | Immunization, measles (% of children ages 12-23 months) | 0.3 (0.22, 0.36) | 98 | Poverty headcount ratio at $1.90 a day (2011 PPP) (% of population) | 0.34 (0.22, 0.36) | 92 |
| 7 | Number of households 5 persons - Per capita | -0.28 (-0.35, -0.1) | 56 | Number of households 6 persons and over - Proportion over All households | -0.29 (-0.37, -0.18) | 53 | Poverty gap at $5.50 a day (2011 PPP) (% of population) | 0.33 (0.23, 0.37) | 92 |

**S5 Results** – Correlation between *R0* and the indicators

***Table S6*** (continued)

| Rank | Pearson | | | Spearman | | | MIC | | |
|---|---|---|---|---|---|---|---|---|---|
| | **Indicator** | **CC** | **N** | **Indicator** | **CC** | **N** | **Indicator** | **CC** | **N** |
| 8 | Proportion of the population aged 0-14 | -0.27 (-0.39, -0.14) | 98 | Health expenditure, total (% of GDP) | 0.28 (0.21, 0.35) | 98 | Poverty headcount ratio at $3.20 a day (2011 PPP) (% of population) | 0.33 (0.22, 0.37) | 92 |
| 9 | Population living in slums (% of urban population) | -0.27 (-0.32, -0.13) | 75 | Urban population (% of total) | 0.28 (0.2, 0.34) | 98 | Proportion of the population aged 0-4 | 0.32 (0.27, 0.38) | 98 |
| 10 | Number of households 6 persons and over - Proportion over All households | -0.26 (-0.34, -0.14) | 53 | Educational attainment, at least completed upper secondary, population 25+, total (%) (cumulative) | 0.27 (0.2, 0.35) | 88 | Physicians (per 1,000 people) | 0.31 (0.25, 0.37) | 98 |
| 11 | Number of households 6 persons and over - Per capita | -0.26 (-0.34, -0.09) | 53 | Immunization, DPT (% of children ages 12-23 months) | 0.26 (0.2, 0.32) | 98 | GDP per capita, PPP (constant 2011 international $) | 0.31 (0.2, 0.33) | 98 |
| 12 | Proportion of the population aged 0-4 | -0.26 (-0.37, -0.13) | 98 | Low-birthweight babies (% of births) | -0.26 (-0.32, -0.19) | 98 | GDP per capita, PPP (current international $) | 0.31 (0.2, 0.33) | 98 |
| 13 | Number of households 2 persons - Proportion over All households | 0.25 (0.13, 0.36) | 56 | Educational attainment, at least completed upper secondary, population 25+, female (%) (cumulative) | 0.26 (0.18, 0.33) | 88 | Probability of dying before age 5 (per 1000 live births) | 0.31 (0.21, 0.33) | 98 |
| 14 | Poverty headcount ratio at $5.50 a day (2011 PPP) (% of population) | -0.25 (-0.33, -0.14) | 92 | Number of households 2 persons - Proportion over All households | 0.25 (0.15, 0.32) | 56 | Low-birthweight babies (% of births) | 0.3 (0.23, 0.39) | 98 |
| 15 | Number of households 1 person - Proportion over All households | 0.25 (0.12, 0.33) | 56 | Educational attainment, at least completed upper secondary, population 25+, male (%) (cumulative) | 0.23 (0.17, 0.31) | 88 | Poverty headcount ratio at national poverty lines (% of population) | 0.3 (0.19, 0.32) | 75 |

**S5 Results** – Correlation between *R0* and the indicators

***Table S6*** (continued)

| Rank | Pearson | | | Spearman | | | MIC | | |
|---|---|---|---|---|---|---|---|---|---|
| | **Indicator** | **CC** | **N** | **Indicator** | **CC** | **N** | **Indicator** | **CC** | **N** |
| 16 | Poverty gap at $5.50 a day (2011 PPP) (% of population) | -0.24 (-0.3, -0.14) | 92 | Proportion of the population aged 0-4 | -0.23 (-0.29, -0.15) | 98 | HDI | 0.29 (0.23, 0.33) | 98 |
| 17 | Poverty headcount ratio at $3.20 a day (2011 PPP) (% of population) | -0.24 (-0.29, -0.14) | 92 | Lifetime risk of maternal death (1 in: rate varies by country) | 0.23 (0.14, 0.29) | 98 | Prevalence of underweight, weight for age (% of children under 5) | 0.29 (0.21, 0.34) | 94 |
| 18 | Urban population (% of total) | 0.22 (0.11, 0.29) | 98 | Proportion of the population aged 0-14 | -0.22 (-0.28, -0.14) | 98 | GDP per capita (1990 Int. GK$) | 0.29 (0.2, 0.32) | 97 |
| 19 | Immunization, measles (% of children ages 12-23 months) | 0.22 (0.03, 0.27) | 98 | Poverty headcount ratio at $3.20 a day (2011 PPP) (% of population) | -0.22 (-0.27, -0.13) | 92 | Number of doctors' consultations | 0.29 (0.2, 0.45) | 25 |
| 20 | Poverty headcount ratio at national poverty lines (% of population) | -0.22 (-0.27, -0.11) | 75 | Proportion of the population aged 65+ | 0.22 (0.15, 0.29) | 98 | Health expenditure, total (% of GDP) | 0.28 (0.21, 0.35) | 98 |
| 21 | Poverty gap at $3.20 a day (2011 PPP) (% of population) | -0.22 (-0.26, -0.13) | 92 | Poverty gap at $5.50 a day (2011 PPP) (% of population) | -0.22 (-0.27, -0.12) | 92 | Number of households 5 persons - Per capita | 0.28 (0.25, 0.42) | 56 |
| 22 | Number of households 1 person - Per capita | 0.21 (0.1, 0.29) | 56 | Poverty headcount ratio at $5.50 a day (2011 PPP) (% of population) | -0.22 (-0.27, -0.12) | 92 | Population growth rate (Average annual rate of population change (percentage)) | 0.27 (0.22, 0.35) | 98 |
| 23 | Poverty headcount ratio at $1.90 a day (2011 PPP) (% of population) | -0.21 (-0.25, -0.12) | 92 | Poverty gap at $3.20 a day (2011 PPP) (% of population) | -0.22 (-0.26, -0.12) | 92 | Immunization, measles (% of children ages 12-23 months) | 0.27 (0.19, 0.32) | 98 |

**S5 Results** – Correlation between *R0* and the indicators

***Table S6*** (continued)

| Rank | Pearson | | | Spearman | | | MIC | | |
|---|---|---|---|---|---|---|---|---|---|
| | **Indicator** | **CC** | **N** | **Indicator** | **CC** | **N** | **Indicator** | **CC** | **N** |
| 24 | Number of households 2 persons - Per capita | 0.21 (0.11 0.3) | 56 | Number of households 4 persons - Proportion over All households | -0.21 (-0.28 -0.11) | 56 | Urban population (% of total) | 0.27 (0.22 0.32) | 98 |
| 25 | Prevalence of underweight, weight for age (% of children under 5) | -0.21 (-0.26 -0.08) | 94 | Exclusive breastfeeding (% of children under 6 months) | 0.21 (0.11 0.27) | 83 | Number of households 3 persons - Per capita | 0.27 (0.22 0.37) | 56 |
| 26 | Population growth rate (Average annual rate of population change (percentage)) | -0.2 (-0.31 -0.13) | 98 | Poverty headcount ratio at $1.90 a day (2011 PPP) (% of population) | -0.21 (-0.26 -0.12) | 92 | Educational attainment, at least completed upper secondary, population 25+, total (%) (cumulative) | 0.27 (0.22 0.36) | 88 |
| 27 | Lifetime risk of maternal death (1 in: rate varies by country) | 0.2 (0.09 0.33) | 98 | HiB vaccination coverage | 0.21 (0.14 0.24) | 96 | Mean age of child-bearing | 0.27 (0.22 0.35) | 98 |
| 28 | HDI | 0.2 (0.08 0.27) | 98 | Immunization, HepB3 (% of one-year-old children) | 0.2 (0.13 0.25) | 91 | Lifetime risk of maternal death (1 in: rate varies by country) | 0.27 (0.22 0.36) | 98 |
| 29 | Immunization, DPT (% of children ages 12-23 months) | 0.19 (0.03 0.25) | 98 | Life expectancy at birth (both sexes) | 0.19 (0.11 0.24) | 98 | Number of households All households - Per capita | 0.27 (0.2 0.34) | 61 |
| 30 | Health expenditure, total (% of GDP) | 0.19 (0.09 0.26) | 98 | Number of households 2 persons - Per capita | 0.19 (0.09 0.27) | 56 | Employment to population ratio, 15+, female (%) (modeled ILO estimate) | 0.26 (0.2 0.3) | 98 |
| 31 | GDP per capita (1990 Int. GK$) | 0.19 (0.09 0.37) | 97 | Poverty headcount ratio at national poverty lines (% of population) | -0.19 (-0.23 -0.08) | 75 | Number of households 2 persons - Proportion over All households | 0.26 (0.21 0.41) | 56 |

**S5 Results** – Correlation between *R0* and the indicators

***Table S6*** (continued)

| Rank | Pearson | | | Spearman | | | MIC | | |
|---|---|---|---|---|---|---|---|---|---|
| | **Indicator** | **CC** | **N** | **Indicator** | **CC** | **N** | **Indicator** | **CC** | **N** |
| 32 | Poverty gap at $1.90 a day (2011 PPP) (%) | -0.18 (-0.22 -0.1) | 92 | GDP per capita (1990 Int. GK$) | 0.19 (0.1 0.24) | 97 | Number of households 5 persons - Proportion over All households | 0.26 (0.23 0.38) | 56 |
| 33 | Exclusive breastfeeding (% of children under 6 months) | 0.18 (0.04 0.23) | 83 | Probability of dying before age 5 (per 1000 live births) | -0.18 (-0.23 -0.1) | 98 | Number of households 4 persons - Proportion over All households | 0.26 (0.21 0.37) | 56 |
| 34 | HiB vaccination coverage | 0.18 (0.06 0.22) | 96 | Total fertility rate (live births per woman) | -0.18 (-0.24 -0.11) | 98 | Number of households 2 persons - Per capita | 0.26 (0.23 0.38) | 56 |
| 35 | Income share held by lowest 10% | 0.18 (0.03 0.3) | 92 | GDP per capita, current prices (Purchasing power parity; international dollars per capita) | 0.18 (0.09 0.22) | 98 | Number of households 6 persons and over - Per capita | 0.25 (0.21 0.41) | 53 |
| 36 | Income share held by lowest 20% | 0.17 (0.03 0.3) | 92 | Prevalence of undernourishment (% of population) | -0.17 (-0.23 -0.06) | 77 | Immunization, DPT (% of children ages 12-23 months) | 0.25 (0.21 0.35) | 98 |
| 37 | Total fertility rate (live births per woman) | -0.17 (-0.28 -0.06) | 98 | Population living in slums (% of urban population) | -0.17 (-0.21 -0.09) | 75 | GDP per capita, current prices (Purchasing power parity; international dollars per capita) | 0.25 (0.2 0.32) | 98 |
| 38 | GDP per capita, current prices (Purchasing power parity; international dollars per capita) | 0.17 (0.06 0.23) | 98 | Number of households 1 person - Proportion over All households | 0.17 (0.08 0.24) | 56 | Immunization, HepB3 (% of one-year-old children) | 0.25 (0.18 0.3) | 91 |
| 39 | Adjusted net enrollment rate, primary (% of primary school age children) | 0.16 (0.05 0.22) | 93 | Population density (people per sq. km of land area) | -0.17 (-0.22 -0.09) | 98 | Total fertility rate (live births per woman) | 0.25 (0.21 0.35) | 98 |

**S5 Results** – Correlation between *R0* and the indicators

***Table S6*** (continued)

| Rank | Pearson | | | Spearman | | | MIC | | |
|---|---|---|---|---|---|---|---|---|---|
| | **Indicator** | **CC** | **N** | **Indicator** | **CC** | **N** | **Indicator** | **CC** | **N** |
| 40 | Mean age of child-bearing | -0.15 (-0.24 -0.04) | 98 | HDI | 0.16 (0.08 0.21) | 98 | Number of households 3 persons - Proportion over All households | 0.25 (0.19 0.35) | 56 |
| 41 | Total population in a household, both sexes | -0.14 (-0.19 -0.01) | 26 | Poverty gap at $1.90 a day (2011 PPP) (%) | -0.15 (-0.21 -0.07) | 92 | Prevalence of undernourishment (% of population) | 0.25 (0.2 0.34) | 77 |
| 42 | Income share held by highest 20% | -0.14 (-0.27 -0.0) | 92 | Adjusted net enrollment rate, primary (% of primary school age children) | 0.15 (0.06 0.2) | 93 | Educational attainment, at least completed upper secondary, population 25+, male (%) (cumulative) | 0.25 (0.21 0.34) | 88 |
| 43 | Life expectancy at birth (both sexes) | 0.13 (0.02 0.23) | 98 | Number of households 1 person - Per capita | 0.15 (0.06 0.23) | 56 | Number of households 6 persons and over - Proportion over All households | 0.24 (0.21 0.42) | 53 |
| 44 | Low-birthweight babies (% of births) | -0.13 (-0.2 -0.06) | 98 | Number of households 4 persons - Per capita | -0.14 (-0.21 -0.03) | 56 | HiB vaccination coverage | 0.24 (0.19 0.3) | 96 |
| 45 | Number of households 3 persons - Per capita | 0.13 (0.04 0.22) | 56 | GDP per capita, PPP (constant 2011 international $) | 0.14 (0.06 0.18) | 98 | Employment to population ratio, ages 15-24, female (%) (modeled ILO estimate) | 0.24 (0.19 0.29) | 98 |
| 46 | Population density (people per sq. km of land area) | -0.1 (-0.12 -0.03) | 98 | Population growth rate (Average annual rate of population change (percentage)) | -0.13 (-0.19 -0.07) | 98 | Unemployment, total (% of total labor force) (modeled ILO estimate) | 0.24 (0.19 0.31) | 98 |
| 47 | GDP per capita, PPP (constant 2011 international $) | 0.09 (0.02 0.19) | 98 | Total population in a household, both sexes | -0.12 (-0.2 -0.01) | 26 | Employment to population ratio, 15+, female (%) (national estimate) | 0.24 (0.19 0.3) | 85 |
| 48 | Number of households All households - Per capita | 0.09 (0.04 0.18) | 61 | GDP per capita, PPP (current international $) | 0.12 (0.05 0.17) | 98 | Number of households 4 persons - Per capita | 0.24 (0.22 0.36) | 56 |

| | | | | | | | | | |
|---|---|---|---|---|---|---|---|---|---|
| 49 | GDP per capita, PPP (current international $) | 0.07 (0.01 0.15) | 98 | Income share held by highest 20% | -0.11 (-0.18 -0.04) | 92 | Proportion of the population aged 0-14 | 0.24 (0.22 0.35) | 98 |

**S5 Results** – Correlation between *R0* and the indicators

***Table S6*** (continued)

| Rank | Pearson | | | Spearman | | | MIC | | |
|---|---|---|---|---|---|---|---|---|---|
| | **Indicator** | **CC** | **N** | **Indicator** | **CC** | **N** | **Indicator** | **CC** | **N** |
| 50 | Average number of people per room in occupied housing unit | 0.48 (-0.21 0.53) | 12 | Income share held by highest 10% | -0.11 (-0.18 -0.04) | 92 | Population living in slums (% of urban population) | 0.24 (0.21 0.34) | 75 |
| 51 | Prevalence of undernourishment (% of population) | -0.15 (-0.2 0.03) | 77 | Employment to population ratio, ages 15-24, female (%) (modeled ILO estimate) | 0.1 (0.04 0.16) | 98 | Population density (people per sq. km of land area) | 0.24 (0.21 0.3) | 98 |
| 52 | Immunization, HepB3 (% of one-year-old children) | 0.15 (-0.01 0.2) | 91 | Income share held by lowest 20% | 0.1 (0.03 0.17) | 92 | Proportion of the population aged 65+ | 0.24 (0.21 0.32) | 98 |
| 53 | Income share held by highest 10% | -0.13 (-0.25 0.01) | 92 | Income share held by lowest 10% | 0.1 (0.03 0.17) | 92 | Number of households 1 person - Per capita | 0.23 (0.2 0.37) | 56 |
| 54 | Probability of dying before age 5 (per 1000 live births) | -0.11 (-0.21 0.02) | 98 | Employment to population ratio, 15+, female (%) (national estimate) | 0.08 (0.01 0.14) | 85 | Exclusive breastfeeding (% of children under 6 months) | 0.23 (0.2 0.34) | 83 |
| 55 | Employment to population ratio, 15+, female (%) (modeled ILO estimate) | -0.11 (-0.14 0.04) | 98 | Average number of people per room in occupied housing unit | 0.29 (-0.15 0.36) | 12 | Employment to population ratio, ages 15-24, female (%) (national estimate) | 0.23 (0.18 0.28) | 80 |
| 56 | Unemployment, total (% of total labor force) (modeled ILO estimate) | -0.11 (-0.15 0.02) | 98 | Number of doctors' consultations | -0.1 (-0.18 0.04) | 25 | Educational attainment, at least completed upper secondary, population 25+, female (%) (cumulative) | 0.23 (0.23 0.36) | 88 |
| 57 | Number of households 4 persons - Proportion over All households | -0.09 (-0.2 0.04) | 56 | Unemployment, total (% of total labor force) (national estimate) | 0.06 (-0.01 0.12) | 98 | Poverty gap at national poverty lines (%) | 0.23 (0.22 0.4) | 51 |

**S5 Results** – Correlation between *R0* and the indicators

***Table S6*** (continued)

| Rank | Pearson | | | Spearman | | | MIC | | |
|---|---|---|---|---|---|---|---|---|---|
| | **Indicator** | **CC** | **N** | **Indicator** | **CC** | **N** | **Indicator** | **CC** | **N** |
| 58 | Number of households 3 persons - Proportion over All households | 0.08 (-0.03 0.18) | 56 | Number of households 3 persons - Per capita | 0.05 (-0.04 0.15) | 56 | Income share held by highest 10% | 0.22 (0.17 0.28) | 92 |
| 59 | Number of doctors' consultations | 0.07 (-0.11 0.25) | 25 | Number of households 3 persons - Proportion over All households | 0.04 (-0.04 0.14) | 56 | Number of households 1 person - Proportion over All households | 0.22 (0.17 0.32) | 56 |
| 60 | Poverty gap at national poverty lines (%) | -0.07 (-0.15 0.04) | 51 | Employment to population ratio, 15+, female (%) (modeled ILO estimate) | -0.03 (-0.08 0.05) | 98 | Adjusted net enrollment rate, primary (% of primary school age children) | 0.22 (0.19 0.33) | 93 |
| 61 | Number of households 4 persons - Per capita | 0.06 (-0.01 0.17) | 56 | Unemployment, total (% of total labor force) (modeled ILO estimate) | 0.02 (-0.05 0.09) | 98 | Unemployment, total (% of total labor force) (national estimate) | 0.21 (0.18 0.27) | 98 |
| 62 | Crude death rate per 1000 population | 0.05 (-0.01 0.14) | 98 | Crude death rate per 1000 population | -0.02 (-0.04 0.07) | 98 | Income share held by lowest 10% | 0.2 (0.18 0.3) | 92 |
| 63 | Employment to population ratio, 15+, female (%) (national estimate) | -0.05 (-0.09 0.09) | 85 | Mean age of child-bearing | -0.02 (-0.08 0.05) | 98 | Income share held by lowest 20% | 0.19 (0.18 0.29) | 92 |
| 64 | Employment to population ratio, ages 15-24, female (%) (national estimate) | -0.03 (-0.07 0.13) | 80 | Poverty gap at national poverty lines (%) | 0.02 (-0.07 0.11) | 51 | Income share held by highest 20% | 0.19 (0.17 0.28) | 92 |
| 65 | Unemployment, total (% of total labor force) (national estimate) | 0.02 (-0.07 0.1) | 98 | Employment to population ratio, ages 15-24, female (%) (national estimate) | 0.01 (-0.05 0.08) | 80 | Total population in a household, both sexes | 0.16 (0.16 0.39) | 26 |
| 66 | Employment to population ratio, ages 15-24, female (%) (modeled ILO estimate) | -0.01 (-0.04 0.17) | 98 | Number of households All households - Per capita | 0.0 (-0.07 0.09) | 61 | Average number of people per room in occupied housing unit | 0.09 (0.09 0.2) | 12 |

#### S6 Results – Effect of studies with high $R_0$ values on the performance of simple linear regression and Random Forest prediction

In this section we explain in detail how the MSE variation of simple linear regression on different indicators can be directly traced to particular combinations of missing indicator values and the value of $R_0$ as calculated using seroprevalence data. Consequently, we argue that this variation in the MSE should not be interpreted as increased predictive power of some indicators over others.

More specifically, we show that those indicators that have a missing value for either or both of the high seroprevalence-estimated $R_0$ value studies ('Czech Republic, <1967' and 'Chile (rural), 1967-68' with $R_0$ value equal to 19.97 and 16.53 respectively) are associated with worse MSE performance. In simple terms, the trained model prediction output cannot reach those high $R_0$ values in either of the two regression methods that we consider (simple linear regression and random forest). Hence the indicators which have a missing value for both those studies have an advantage in terms of the prediction MSE.

Contrary to that, that indicators which have a valid value for both those studies are associated with a higher MSE. Furthermore, this increase in the MSE is further exacerbated for those indicators which have very few valid values overall, as the higher error due to the two high $R_0$ studies is averaged over a smaller number of total studies. Hence, indicators with a valid value for the 'Czech Republic, <1967' and 'Chile (rural), 1967-68' studies but with valid values for a few studies overall are associated with the highest MSE.

Starting from the simple linear regression case, as can be seen in Table S7 (which also has a version sorted by MSE value for presentation clarity), the best performing indicator ('Poverty gap at national poverty lines (%)' –listed in line 20 of Table S7 with MSE value equal to 2.73) is the only indicator having a missing value for both those studies (this indicator also has a missing value for the third highest $R_0$ study which is 'East Germany, 1990[45]' with $R_0$ equal to 10.19).

Furthermore, 4 out of the 6 indicators that have a missing value in exactly one of those two studies, are directly following in terms of MSE performance ('Adjusted net enrollment rate, primary (% of primary school age children)', 'Prevalence of undernourishment (% of population)', 'Exclusive breastfeeding (% of children under 6 months)' and 'Population living in slums (% of urban population)' listed in line numbers 30, 41, 47 and 52 of Table S7 with MSE values equal to 5.35, 5.5, 5.36 and 5.39 respectively). Those 4 indicators have valid values for at least 75 studies in total. The remaining two indicators which have a missing value in exactly one of the two high $R_0$ studies ('Average number of people per room in occupied housing unit' and 'Total population in a household, both sexes') have valid values for only 12 and 26 studies in total (see Table S7). As was described above, this practically means that the higher error due to high $R_0$ study is averaged over a much smaller number of total studies and, consequently, the MSE for those two indicators is much higher (33.32 and 17.4 respectively).

Considering the indicators which have a valid value for both of the high $R_0$ studies ('Czech Republic, <1967' and 'Chile (rural), 1967-68') the aforementioned effects result in a clear three-way division in terms of performance: A total of 45 indicators have valid values for 75 up to 98 studies and the MSE associated with them is clustered between the levels of 7.41 and 9.57 (those are the indicators listed in Table S7 in positions 1-19, 21-29, 31-40, 42-46, 48 and 50). A second group contains 13 indicators which have valid values for 53 up to 61 studies and the MSE associated with them is clustered between the levels of 10.15 and 12.26 (those are the indicators listed in Table S7 in positions 53-65). A third group contains one indicator ('Number of doctors' consultations') which has valid values for only 25 studies in total and for which the MSE raises to a value of 26.93

A similar effect can be seen in the random forest performance results regarding the indicator subsets that are used to train and the random forest (a tabular summary is included in the ***Table S8***). In the cross validation experiment conducted using only the 25 indicators that have no missing values, both the high $R_0$ studies are present and the overall error has the value of 9.87. In the experiments conducted using the indicators that have up to 10, 20 and 40 missing values only one of the two high $R_0$ studies is present and the overall error is lower. However, as the number of studies involved in the experiment reduces from 69 to 52 and then 32 as we allow indicators with progressively more missing values, the overall error progressively increases (8.19, 9.6 and 13.8 respectively). Finally, none of the two high $R_0$ studies is included in the experiment conducted with the 20 indicators having up to 70 missing values and the overall error in the case attains the lowest value of 6.88.

None of those effects can be seen in the imputed results (Fig 3B of the main text), with the exception of a small increase in the max range of the error value for the indicators ‘Number of doctors' consultations’ and ‘Average number of people per room in occupied housing unit’ indicators which are the ones with the fewest valid values overall (and hence the most imputed values).

Finally, in **Fig S3** we plot the MSE values calculated in the same way as for Fig 3 of the main text but when the cross-validation experiment is run after excluding the ‘Czech Republic, <1967’ and ‘Chile (rural), 1967-68’ studies. In this case the highest value of $R_0$ as estimated from seroprevalence data is for the ‘East Germany, 1990’ study ($R_0$=10.19). Most of the MSE variation can be seen in **Fig S3** to be lost in this case. The only variation is a small decrease in the MSE value for the 5 indicators which have a missing value for the East Germany study (‘Poverty gap at national poverty lines (%)’, ‘Poverty headcount ratio at national poverty lines (% of population)’, ‘Prevalence of undernourishment (% of population)’, ‘Exclusive breastfeeding (% of children under 6 months)’ and ‘Population living in slums (% of urban population)’) and an increase in the MSE for the indicators with the fewest overall valid values

(‘Average number of people per room in occupied housing unit’ and ‘Number of doctors' consultations’). Both those effects follow exactly the same pattern described above.

**Table S7:** Presence or absence of the two studies with highest value of $R_0$ from non-imputed linear regression 4-fold cross validation experiments

| | | Chile (rural), 1967-68 | Czech Republic, <1967 | Average MSE over 10 repetitions | Number of studies with valid values |
|---|---|---|---|---|---|
| 1 | Proportion of the population aged 0-4 | + | + | 7.81 | 98 |
| 2 | Proportion of the population aged 0-14 | + | + | 7.77 | 98 |
| 3 | Proportion of the population aged 65+ | + | + | 7.51 | 98 |
| 4 | Lifetime risk of maternal death (1 in: rate varies by country) | + | + | 7.96 | 98 |
| 5 | Probability of dying before age 5 (per 1000 live births) | + | + | 8.12 | 98 |
| 6 | Crude death rate per 1000 population | + | + | 8.06 | 98 |
| 7 | Life expectancy at birth (both sexes) | + | + | 8.04 | 98 |
| 8 | Total fertility rate (live births per woman) | + | + | 8.04 | 98 |
| 9 | Mean age of child-bearing | + | + | 8.32 | 98 |
| 10 | Population growth rate (Average annual rate of population change (percentage)) | + | + | 7.98 | 98 |
| 11 | Population density (people per sq. km of land area) | + | + | 8.11 | 98 |
| 12 | Urban population (% of total) | + | + | 7.76 | 98 |
| 13 | Income share held by highest 10% | + | + | 9.21 | 92 |
| 14 | Income share held by highest 20% | + | + | 9.2 | 92 |
| 15 | Income share held by lowest 10% | + | + | 9.08 | 92 |
| 16 | Income share held by lowest 20% | + | + | 9.03 | 92 |
| 17 | Poverty gap at $1.90 a day (2011 PPP) (%) | + | + | 8.27 | 92 |
| 18 | Poverty gap at $3.20 a day (2011 PPP) (% of population) | + | + | 8.22 | 92 |
| 19 | Poverty gap at $5.50 a day (2011 PPP) (% of population) | + | + | 8.19 | 92 |
| 20 | Poverty gap at national poverty lines (%) | - | - | 2.73 | 51 |
| 21 | Poverty headcount ratio at $1.90 a day (2011 PPP) (% of population) | + | + | 8.24 | 92 |
| 22 | Poverty headcount ratio at $3.20 a day (2011 PPP) (% of population) | + | + | 8.21 | 92 |
| 23 | Poverty headcount ratio at $5.50 a day (2011 PPP) (% of population) | + | + | 8.24 | 92 |
| 24 | Poverty headcount ratio at national poverty lines (% of population) | + | + | 9.46 | 75 |
| 25 | GDP per capita, PPP (constant 2011 international $) | + | + | 8.06 | 98 |
| 26 | GDP per capita, PPP (current international $) | + | + | 8.09 | 98 |
| 27 | HDI | + | + | 7.84 | 98 |
| 28 | GDP per capita, current prices (Purchasing power parity; international dollars per capita) | + | + | 8.07 | 98 |
| 29 | GDP per capita (1990 Int. GK$) | + | + | 7.89 | 97 |
| 30 | Adjusted net enrollment rate, primary (% of primary school age children) | + | - | 5.35 | 93 |
| 31 | Educational attainment, at least completed upper secondary, population 25+, female (%) (cumulative) | + | + | 7.72 | 88 |

| | | | | | |
|---|---|---|---|---|---|
| 32 | Educational attainment, at least completed upper secondary, population 25+, male (%) (cumulative) | + | + | 7.76 | 88 |
| 33 | Educational attainment, at least completed upper secondary, population 25+, total (%) (cumulative) | + | + | 7.69 | 88 |
| 34 | Unemployment, total (% of total labor force) (modeled ILO estimate) | + | + | 8.1 | 98 |
| 35 | Unemployment, total (% of total labor force) (national estimate) | + | + | 8.26 | 98 |
| 36 | Employment to population ratio, 15+, female (%) (modeled ILO estimate) | + | + | 8 | 98 |
| 37 | Employment to population ratio, 15+, female (%) (national estimate) | + | + | 9.04 | 85 |
| 38 | Employment to population ratio, ages 15-24, female (%) (modeled ILO estimate) | + | + | 8.11 | 98 |
| 39 | Employment to population ratio, ages 15-24, female (%) (national estimate) | + | + | 9.57 | 80 |
| 40 | Low-birthweight babies (% of births) | + | + | 8.06 | 98 |
| 41 | Prevalence of undernourishment (% of population) | + | - | 5.5 | 77 |
| 42 | Prevalence of underweight, weight for age (% of children under 5) | + | + | 8.22 | 94 |
| 43 | Health expenditure, total (% of GDP) | + | + | 7.83 | 98 |
| 44 | Immunization, DPT (% of children ages 12-23 months) | + | + | 8 | 98 |
| 45 | Immunization, HepB3 (% of one-year-old children) | + | + | 8.57 | 91 |
| 46 | Immunization, measles (% of children ages 12-23 months) | + | + | 7.91 | 98 |
| 47 | Exclusive breastfeeding (% of children under 6 months) | + | - | 5.36 | 83 |
| 48 | Physicians (per 1,000 people) | + | + | 7.41 | 98 |
| 49 | Number of doctors' consultations | + | + | 26.21 | 25 |
| 50 | HiB vaccination coverage | + | + | 8.07 | 96 |
| 51 | Average number of people per room in occupied housing unit | - | + | 33.32 | 12 |
| 52 | Population living in slums (% of urban population) | + | - | 5.39 | 75 |
| 53 | Number of households All households - Per capita | + | + | 11.51 | 61 |
| 54 | Number of households 1 person - Per capita | + | + | 11.82 | 56 |
| 55 | Number of households 1 person - Proportion over All households | + | + | 11.89 | 56 |
| 56 | Number of households 2 persons - Per capita | + | + | 11.66 | 56 |
| 57 | Number of households 2 persons - Proportion over All households | + | + | 11.59 | 56 |
| 58 | Number of households 3 persons - Per capita | + | + | 11.94 | 56 |
| 59 | Number of households 3 persons - Proportion over All households | + | + | 12.09 | 56 |
| 60 | Number of households 4 persons - Per capita | + | + | 12.19 | 56 |
| 61 | Number of households 4 persons - Proportion over All households | + | + | 12.26 | 56 |
| 62 | Number of households 5 persons - Per capita | + | + | 11.64 | 56 |
| 63 | Number of households 5 persons - Proportion over All households | + | + | 11.15 | 56 |
| 64 | Number of households 6 persons and over - Per capita | + | + | 12.07 | 53 |
| 65 | Number of households 6 persons and over - Proportion over All households | + | + | 11.96 | 53 |
| 66 | Total population in a household, both sexes | - | + | 17.4 | 26 |

**Table S7 (sorted by Average MSE value):** Presence or absence of the two studies with highest value of $R_0$ from non-imputed linear regression 4-fold cross validation experiments

| | | Chile (rural), 1967-68 | Czech Republic, <1967 | Average MSE over 10 repetitions | Number of studies with valid values |
|---|---|---|---|---|---|
| 20 | Poverty gap at national poverty lines (%) | - | - | 2.73 | 51 |
| 30 | Adjusted net enrollment rate, primary (% of primary school age children) | + | - | 5.35 | 93 |
| 47 | Exclusive breastfeeding (% of children under 6 months) | + | - | 5.36 | 83 |
| 52 | Population living in slums (% of urban population) | + | - | 5.39 | 75 |
| 41 | Prevalence of undernourishment (% of population) | + | - | 5.5 | 77 |
| 48 | Physicians (per 1,000 people) | + | + | 7.41 | 98 |
| 3 | Proportion of the population aged 65+ | + | + | 7.51 | 98 |
| 33 | Educational attainment, at least completed upper secondary, population 25+, total (%) (cumulative) | + | + | 7.69 | 88 |
| 31 | Educational attainment, at least completed upper secondary, population 25+, female (%) (cumulative) | + | + | 7.72 | 88 |
| 12 | Urban population (% of total) | + | + | 7.76 | 98 |
| 32 | Educational attainment, at least completed upper secondary, population 25+, male (%) (cumulative) | + | + | 7.76 | 88 |
| 2 | Proportion of the population aged 0-14 | + | + | 7.77 | 98 |
| 1 | Proportion of the population aged 0-4 | + | + | 7.81 | 98 |
| 43 | Health expenditure, total (% of GDP) | + | + | 7.83 | 98 |
| 27 | HDI | + | + | 7.84 | 98 |
| 29 | GDP per capita (1990 Int. GK$) | + | + | 7.89 | 97 |
| 46 | Immunization, measles (% of children ages 12-23 months) | + | + | 7.91 | 98 |
| 4 | Lifetime risk of maternal death (1 in: rate varies by country) | + | + | 7.96 | 98 |
| 10 | Population growth rate (Average annual rate of population change (percentage)) | + | + | 7.98 | 98 |
| 36 | Employment to population ratio, 15+, female (%) (modeled ILO estimate) | + | + | 8 | 98 |
| 44 | Immunization, DPT (% of children ages 12-23 months) | + | + | 8 | 98 |
| 7 | Life expectancy at birth (both sexes) | + | + | 8.04 | 98 |
| 8 | Total fertility rate (live births per woman) | + | + | 8.04 | 98 |
| 6 | Crude death rate per 1000 population | + | + | 8.06 | 98 |
| 25 | GDP per capita, PPP (constant 2011 international $) | + | + | 8.06 | 98 |
| 40 | Low-birthweight babies (% of births) | + | + | 8.06 | 98 |
| 28 | GDP per capita, current prices (Purchasing power parity; international dollars per capita) | + | + | 8.07 | 98 |
| 50 | HiB vaccination coverage | + | + | 8.07 | 96 |
| 26 | GDP per capita, PPP (current international $) | + | + | 8.09 | 98 |
| 34 | Unemployment, total (% of total labor force) (modeled ILO estimate) | + | + | 8.1 | 98 |
| 11 | Population density (people per sq. km of land area) | + | + | 8.11 | 98 |
| 38 | Employment to population ratio, ages 15-24, female (%) (modeled ILO estimate) | + | + | 8.11 | 98 |

| | | | | | |
|---|---|---|---|---|---|
| 5 | Probability of dying before age 5 (per 1000 live births) | + | + | 8.12 | 98 |
| 19 | Poverty gap at $5.50 a day (2011 PPP) (% of population) | + | + | 8.19 | 92 |
| 22 | Poverty headcount ratio at $3.20 a day (2011 PPP) (% of population) | + | + | 8.21 | 92 |
| 18 | Poverty gap at $3.20 a day (2011 PPP) (% of population) | + | + | 8.22 | 92 |
| 42 | Prevalence of underweight, weight for age (% of children under 5) | + | + | 8.22 | 94 |
| 21 | Poverty headcount ratio at $1.90 a day (2011 PPP) (% of population) | + | + | 8.24 | 92 |
| 23 | Poverty headcount ratio at $5.50 a day (2011 PPP) (% of population) | + | + | 8.24 | 92 |
| 35 | Unemployment, total (% of total labor force) (national estimate) | + | + | 8.26 | 98 |
| 17 | Poverty gap at $1.90 a day (2011 PPP) (%) | + | + | 8.27 | 92 |
| 9 | Mean age of child-bearing | + | + | 8.32 | 98 |
| 45 | Immunization, HepB3 (% of one-year-old children) | + | + | 8.57 | 91 |
| 16 | Income share held by lowest 20% | + | + | 9.03 | 92 |
| 37 | Employment to population ratio, 15+, female (%) (national estimate) | + | + | 9.04 | 85 |
| 15 | Income share held by lowest 10% | + | + | 9.08 | 92 |
| 14 | Income share held by highest 20% | + | + | 9.2 | 92 |
| 13 | Income share held by highest 10% | + | + | 9.21 | 92 |
| 24 | Poverty headcount ratio at national poverty lines (% of population) | + | + | 9.46 | 75 |
| 39 | Employment to population ratio, ages 15-24, female (%) (national estimate) | + | + | 9.57 | 80 |
| 63 | Number of households 5 persons - Proportion over All households | + | + | 11.15 | 56 |
| 53 | Number of households All households - Per capita | + | + | 11.51 | 61 |
| 57 | Number of households 2 persons - Proportion over All households | + | + | 11.59 | 56 |
| 62 | Number of households 5 persons - Per capita | + | + | 11.64 | 56 |
| 56 | Number of households 2 persons - Per capita | + | + | 11.66 | 56 |
| 54 | Number of households 1 person - Per capita | + | + | 11.82 | 56 |
| 55 | Number of households 1 person - Proportion over All households | + | + | 11.89 | 56 |
| 58 | Number of households 3 persons - Per capita | + | + | 11.94 | 56 |
| 65 | Number of households 6 persons and over - Proportion over All households | + | + | 11.96 | 53 |
| 64 | Number of households 6 persons and over - Per capita | + | + | 12.07 | 53 |
| 59 | Number of households 3 persons - Proportion over All households | + | + | 12.09 | 56 |
| 60 | Number of households 4 persons - Per capita | + | + | 12.19 | 56 |
| 61 | Number of households 4 persons - Proportion over All households | + | + | 12.26 | 56 |
| 66 | Total population in a household, both sexes | - | + | 17.4 | 26 |
| 49 | Number of doctors' consultations | + | + | 26.21 | 25 |
| 51 | Average number of people per room in occupied housing unit | - | + | 33.32 | 12 |

**Table S8:** Presence or absence of the two studies with highest value of $R_0$ from non-imputed random forest 4-fold cross validation experiments

| | Chile (rural), 1967-68 | Czech Republic, <1967 | Average MSE over 10 repetitions | Number of studies with valid values |
|---|---|---|---|---|
| Using all indicators with no missing values | + | + | 9.87 | 98 |
| Using all indicators with up to 10 missing values | + | - | 8.19 | 69 |
| Using all indicators with up to 20 missing values | + | - | 9.6 | 52 |
| Using all indicators with up to 40 missing values | + | - | 13.8 | 32 |
| Using all indicators with up to 70 missing values | - | - | 6.88 | 20 |

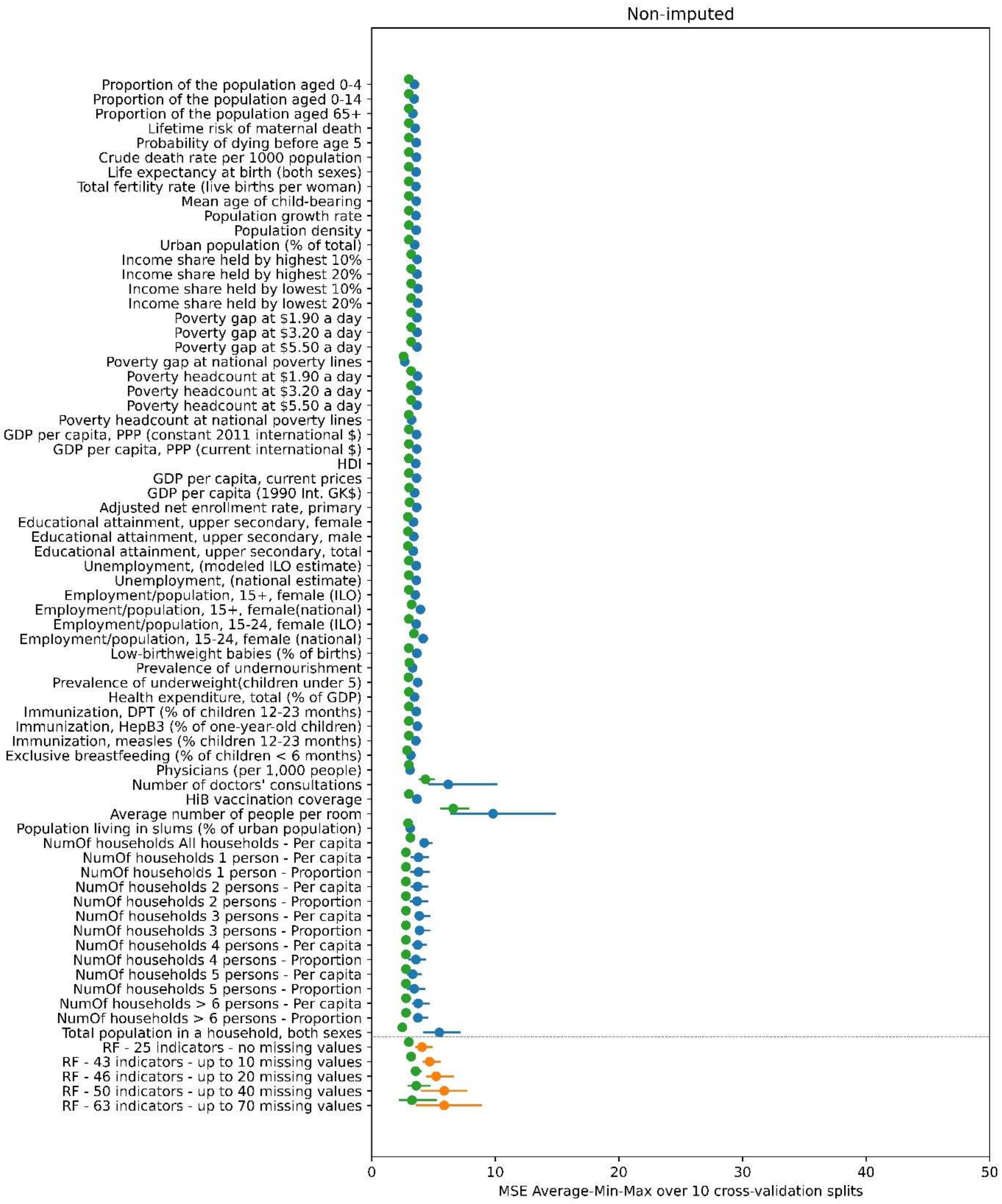

**Fig S3A**: *:* Similar to Fig 3A of the main text but with the two studies ('Czech Republic, <1967' and 'Chile (rural), 1967-68') excluded. Mean value (blue and orange dots) and minimum-maximum value range (blue and orange line) of the MSE of the predicted $R_0$ over ten 4-fold cross validation splits

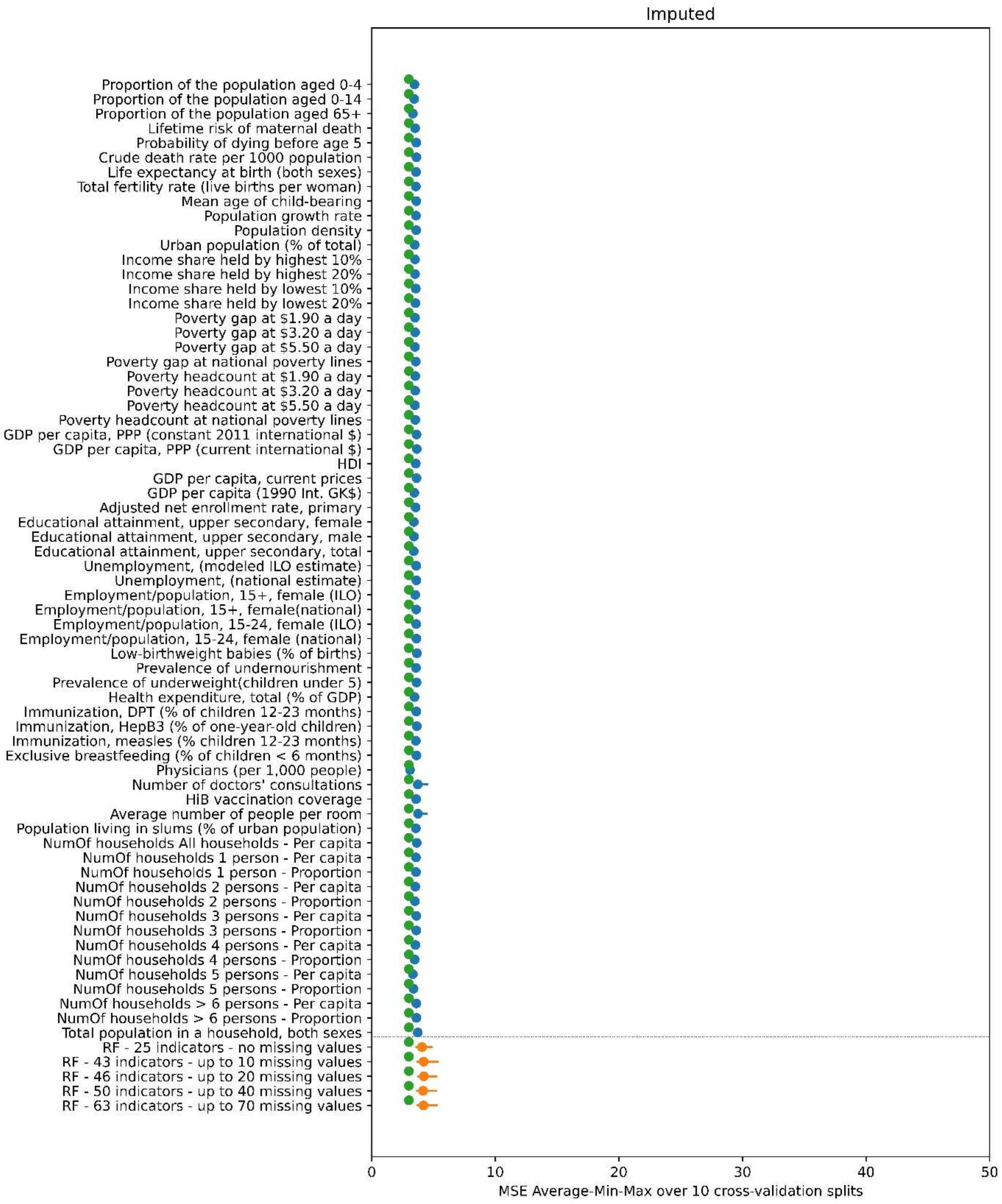

**Fig S3B**: Similar to Fig 3B of the main text but with the two studies ('Czech Republic, <1967' and 'Chile (rural), 1967-68') excluded. Mean value (blue and orange dots) and minimum-maximum value range (blue and orange line) of the MSE of the predicted $R_0$ over ten 4-fold cross validation splits

### S7 Results – Random forest parameter tuning

For the parameter optimisation results we present in the main test we run a nested cross-validation experiment over a grid spanning 6 parameters and a total of 96 points. We chose these parameters to include following the description in the documentation of scikit-learn documentation (https://scikit-learn.org/stable/modules/ensemble.html). The 6 parameters together with their default value (in brackets) and their alternative values are listed below:

- ‘bootstrap’: [True], False
- ‘max_depth’: [None], 10
- ‘max_features’: [‘auto’], ‘sqrt’
- ‘min_samples_leaf’: [1], 2
- ‘min_samples_split’: [2], 4
- ‘n_estimators’: [100], 25, 200

In **Table S9** we list the MSE (average, minimum and maximum over the 10 splits) obtained from the nested cross-validation optimisation run for the 96 points of the parameters grid. We also list the rank (average, minimum and maximum over the 10 splits) in terms of the achieved MSE for each grid point. For the ‘bootstrap’ and ‘max_features’ parameters for which the ‘TRUE and ‘sqrt’ values are performing consistently better. For the other 4 parameters there does not seem to be a clear separation between better and worse performing values.

**Table S9**: MSE performance and rank of the 96 parameter sets in the double-loop nested cross-validation experiment. The default parameter setup is in bold font.

| bootstrap | max_depth | max_features | min_samples_leaf | min_samples_split | n_estimators | Mean MSE OverSplits | Min MSE OverSplits | Max MSE OverSplits | Mean Rank OverSplits | MinRankOverSplits | MaxRankOverSplits |
|---|---|---|---|---|---|---|---|---|---|---|---|
| TRUE | 10 | sqrt | 2 | 2 | 200 | 8.41 | 8.13 | 8.70 | 4.5 | 1 | 11 |
| TRUE | 10 | sqrt | 2 | 4 | 200 | 8.41 | 8.13 | 8.70 | 4.5 | 1 | 11 |
| TRUE | None | sqrt | 2 | 2 | 200 | 8.41 | 8.13 | 8.73 | 4.6 | 1 | 9 |
| TRUE | None | sqrt | 2 | 4 | 200 | 8.41 | 8.13 | 8.73 | 4.6 | 1 | 9 |
| TRUE | None | sqrt | 2 | 2 | 100 | 8.46 | 8.10 | 8.76 | 6 | 1 | 11 |
| TRUE | None | sqrt | 2 | 4 | 100 | 8.46 | 8.10 | 8.76 | 6 | 1 | 11 |
| TRUE | 10 | sqrt | 2 | 2 | 100 | 8.46 | 8.10 | 8.75 | 7.5 | 3 | 12 |
| TRUE | 10 | sqrt | 2 | 4 | 100 | 8.46 | 8.10 | 8.75 | 7.5 | 3 | 12 |
| TRUE | None | sqrt | 2 | 2 | 25 | 8.58 | 8.05 | 9.01 | 11.3 | 1 | 33 |
| TRUE | None | sqrt | 2 | 4 | 25 | 8.58 | 8.05 | 9.01 | 11.3 | 1 | 33 |
| TRUE | 10 | sqrt | 2 | 2 | 25 | 8.60 | 8.05 | 8.92 | 12.1 | 1 | 30 |
| TRUE | 10 | sqrt | 2 | 4 | 25 | 8.60 | 8.05 | 8.92 | 12.1 | 1 | 30 |
| TRUE | None | sqrt | 1 | 4 | 200 | 8.78 | 8.43 | 9.14 | 16.1 | 9 | 26 |
| TRUE | 10 | sqrt | 1 | 4 | 200 | 8.78 | 8.46 | 9.16 | 17.2 | 9 | 28 |
| TRUE | None | sqrt | 1 | 4 | 100 | 8.81 | 8.53 | 9.36 | 17.3 | 13 | 25 |
| TRUE | 10 | sqrt | 1 | 4 | 100 | 8.80 | 8.53 | 9.37 | 17.9 | 10 | 28 |
| TRUE | 10 | auto | 2 | 2 | 200 | 8.99 | 8.41 | 9.64 | 22.5 | 13 | 33 |
| TRUE | 10 | auto | 2 | 4 | 200 | 8.99 | 8.41 | 9.64 | 22.5 | 13 | 33 |
| TRUE | None | auto | 2 | 2 | 200 | 8.99 | 8.41 | 9.64 | 22.9 | 15 | 31 |
| TRUE | None | auto | 2 | 4 | 200 | 8.99 | 8.41 | 9.64 | 22.9 | 15 | 31 |
| TRUE | 10 | auto | 2 | 2 | 100 | 9.04 | 8.52 | 9.84 | 23.7 | 11 | 37 |
| TRUE | 10 | auto | 2 | 4 | 100 | 9.04 | 8.52 | 9.84 | 23.7 | 11 | 37 |
| TRUE | None | auto | 2 | 2 | 100 | 9.04 | 8.54 | 9.85 | 24.4 | 13 | 37 |
| TRUE | None | auto | 2 | 4 | 100 | 9.04 | 8.54 | 9.85 | 24.4 | 13 | 37 |
| TRUE | 10 | sqrt | 1 | 4 | 25 | 9.03 | 8.47 | 9.61 | 25.7 | 9 | 48 |
| TRUE | None | sqrt | 1 | 4 | 25 | 9.02 | 8.49 | 9.63 | 26.3 | 14 | 50 |
| TRUE | None | sqrt | 1 | 2 | 200 | 9.09 | 8.62 | 9.67 | 27 | 17 | 36 |
| TRUE | 10 | sqrt | 1 | 2 | 200 | 9.11 | 8.60 | 9.70 | 27.6 | 14 | 37 |
| TRUE | None | sqrt | 1 | 2 | 100 | 9.12 | 8.62 | 9.85 | 28 | 16 | 37 |
| TRUE | 10 | sqrt | 1 | 2 | 100 | 9.15 | 8.64 | 9.94 | 28.4 | 10 | 38 |
| TRUE | None | auto | 2 | 2 | 25 | 9.28 | 8.69 | 10.51 | 34.3 | 3 | 59 |
| TRUE | None | auto | 2 | 4 | 25 | 9.28 | 8.69 | 10.51 | 34.3 | 3 | 59 |
| TRUE | 10 | auto | 2 | 2 | 25 | 9.28 | 8.65 | 10.49 | 34.6 | 1 | 55 |
| TRUE | 10 | auto | 2 | 4 | 25 | 9.28 | 8.65 | 10.49 | 34.6 | 1 | 55 |
| TRUE | 10 | sqrt | 1 | 2 | 25 | 9.35 | 8.47 | 10.23 | 36.3 | 14 | 47 |
| TRUE | None | sqrt | 1 | 2 | 25 | 9.36 | 8.29 | 10.27 | 37 | 13 | 58 |
| TRUE | 10 | auto | 1 | 4 | 100 | 9.54 | 8.63 | 10.61 | 41 | 18 | 54 |
| TRUE | None | auto | 1 | 4 | 100 | 9.54 | 8.66 | 10.60 | 41.4 | 23 | 53 |
| TRUE | 10 | auto | 1 | 4 | 25 | 9.64 | 8.55 | 11.55 | 41.8 | 15 | 68 |

| | | | | | | | | | | | |
|---|---|---|---|---|---|---|---|---|---|---|---|
| TRUE | None | auto | 1 | 4 | 200 | 9.55 | 8.74 | 10.37 | 41.8 | 15 | 55 |
| TRUE | None | auto | 1 | 4 | 25 | 9.63 | 8.60 | 11.53 | 41.9 | 22 | 67 |
| TRUE | 10 | auto | 1 | 4 | 200 | 9.55 | 8.77 | 10.39 | 42 | 16 | 56 |
| FALSE | None | sqrt | 2 | 2 | 100 | 9.57 | 9.11 | 10.27 | 43.9 | 27 | 54 |
| FALSE | None | sqrt | 2 | 4 | 100 | 9.57 | 9.11 | 10.27 | 43.9 | 27 | 54 |
| FALSE | 10 | sqrt | 2 | 2 | 100 | 9.57 | 9.11 | 10.16 | 44.1 | 25 | 55 |
| FALSE | 10 | sqrt | 2 | 4 | 100 | 9.57 | 9.11 | 10.16 | 44.1 | 25 | 55 |
| FALSE | None | sqrt | 2 | 2 | 200 | 9.62 | 9.10 | 10.33 | 45.4 | 31 | 59 |
| FALSE | None | sqrt | 2 | 4 | 200 | 9.62 | 9.10 | 10.33 | 45.4 | 31 | 59 |
| FALSE | 10 | sqrt | 2 | 2 | 200 | 9.61 | 9.11 | 10.19 | 45.5 | 29 | 60 |
| FALSE | 10 | sqrt | 2 | 4 | 200 | 9.61 | 9.11 | 10.19 | 45.5 | 29 | 60 |
| FALSE | 10 | sqrt | 2 | 2 | 25 | 9.63 | 9.22 | 10.13 | 47.3 | 21 | 62 |
| FALSE | 10 | sqrt | 2 | 4 | 25 | 9.63 | 9.22 | 10.13 | 47.3 | 21 | 62 |
| FALSE | None | sqrt | 2 | 2 | 25 | 9.66 | 9.02 | 10.10 | 47.8 | 27 | 65 |
| FALSE | None | sqrt | 2 | 4 | 25 | 9.66 | 9.02 | 10.10 | 47.8 | 27 | 65 |
| TRUE | None | auto | 1 | 2 | 200 | 9.85 | 8.87 | 10.73 | 50.5 | 27 | 60 |
| TRUE | 10 | auto | 1 | 2 | 200 | 9.85 | 8.92 | 10.76 | 50.7 | 32 | 60 |
| TRUE | 10 | auto | 1 | 2 | 100 | 9.85 | 9.00 | 11.01 | 51.1 | 36 | 60 |
| TRUE | None | auto | 1 | 2 | 100 | 9.87 | 8.95 | 11.11 | 51.4 | 35 | 61 |
| TRUE | None | auto | 1 | 2 | 25 | 10.00 | 8.87 | 12.13 | 52 | 38 | 72 |
| TRUE | 10 | auto | 1 | 2 | 25 | 9.98 | 8.91 | 11.79 | 52.2 | 40 | 71 |
| FALSE | None | sqrt | 1 | 4 | 100 | 10.41 | 9.18 | 11.67 | 59.5 | 49 | 64 |
| FALSE | None | sqrt | 1 | 4 | 200 | 10.42 | 9.21 | 11.58 | 60.1 | 51 | 64 |
| FALSE | 10 | sqrt | 1 | 4 | 200 | 10.44 | 9.25 | 11.46 | 60.6 | 52 | 66 |
| FALSE | 10 | sqrt | 1 | 4 | 100 | 10.46 | 9.27 | 11.53 | 61.1 | 53 | 67 |
| FALSE | None | sqrt | 1 | 4 | 25 | 10.65 | 9.40 | 11.75 | 62.9 | 59 | 66 |
| FALSE | 10 | sqrt | 1 | 4 | 25 | 10.78 | 9.46 | 12.21 | 64.3 | 61 | 66 |
| FALSE | None | sqrt | 1 | 2 | 100 | 11.18 | 9.84 | 12.58 | 69.3 | 64 | 82 |
| FALSE | None | sqrt | 1 | 2 | 200 | 11.22 | 9.79 | 12.62 | 69.5 | 63 | 83 |
| FALSE | 10 | sqrt | 1 | 2 | 200 | 11.22 | 9.76 | 12.66 | 69.6 | 65 | 81 |
| FALSE | 10 | sqrt | 1 | 2 | 100 | 11.22 | 9.81 | 12.87 | 69.8 | 66 | 80 |
| FALSE | None | sqrt | 1 | 2 | 25 | 11.46 | 9.99 | 13.04 | 71.8 | 67 | 84 |
| FALSE | 10 | sqrt | 1 | 2 | 25 | 11.46 | 9.84 | 13.27 | 72.7 | 65 | 84 |
| FALSE | 10 | auto | 2 | 2 | 25 | 13.16 | 11.36 | 18.22 | 80.8 | 71 | 91 |
| FALSE | 10 | auto | 2 | 4 | 25 | 13.16 | 11.36 | 18.22 | 80.8 | 71 | 91 |
| FALSE | None | auto | 2 | 2 | 25 | 13.14 | 11.42 | 18.00 | 80.8 | 71 | 93 |
| FALSE | None | auto | 2 | 2 | 100 | 13.18 | 11.38 | 18.27 | 80.8 | 73 | 89 |
| FALSE | None | auto | 2 | 4 | 25 | 13.14 | 11.42 | 18.00 | 80.8 | 71 | 93 |
| FALSE | None | auto | 2 | 4 | 100 | 13.18 | 11.38 | 18.27 | 80.8 | 73 | 89 |
| FALSE | 10 | auto | 2 | 2 | 100 | 13.18 | 11.29 | 18.32 | 81.4 | 73 | 95 |
| FALSE | 10 | auto | 2 | 4 | 100 | 13.18 | 11.29 | 18.32 | 81.4 | 73 | 95 |
| FALSE | 10 | auto | 2 | 2 | 200 | 13.19 | 11.24 | 18.41 | 81.6 | 67 | 95 |
| FALSE | 10 | auto | 2 | 4 | 200 | 13.19 | 11.24 | 18.41 | 81.6 | 67 | 95 |
| FALSE | None | auto | 2 | 2 | 200 | 13.19 | 11.30 | 18.42 | 81.6 | 69 | 95 |

| | | | | | | | | | | | |
|---|---|---|---|---|---|---|---|---|---|---|---|
| FALSE | None | auto | 2 | 4 | 200 | 13.19 | 11.30 | 18.42 | 81.6 | 69 | 95 |
| FALSE | 10 | auto | 1 | 4 | 25 | 14.99 | 10.81 | 19.30 | 81.8 | 73 | 90 |
| FALSE | None | auto | 1 | 4 | 25 | 14.97 | 10.65 | 19.06 | 82.7 | 73 | 89 |
| FALSE | None | auto | 1 | 4 | 200 | 15.02 | 10.61 | 19.07 | 83.6 | 73 | 89 |
| FALSE | 10 | auto | 1 | 4 | 200 | 15.05 | 10.71 | 19.24 | 83.8 | 76 | 90 |
| FALSE | 10 | auto | 1 | 4 | 100 | 15.07 | 10.69 | 19.27 | 84 | 76 | 90 |
| FALSE | None | auto | 1 | 4 | 100 | 15.03 | 10.62 | 19.05 | 84.1 | 74 | 90 |
| FALSE | 10 | auto | 1 | 2 | 25 | 16.23 | 11.16 | 21.21 | 89.1 | 80 | 96 |
| FALSE | None | auto | 1 | 2 | 25 | 16.26 | 11.22 | 21.05 | 89.3 | 79 | 96 |
| FALSE | None | auto | 1 | 2 | 200 | 16.27 | 11.10 | 21.17 | 89.8 | 79 | 95 |
| FALSE | None | auto | 1 | 2 | 100 | 16.29 | 11.10 | 21.18 | 90.1 | 79 | 96 |
| FALSE | 10 | auto | 1 | 2 | 200 | 16.31 | 11.22 | 21.38 | 90.2 | 81 | 96 |
| FALSE | 10 | auto | 1 | 2 | 100 | 16.33 | 11.25 | 21.43 | 90.9 | 83 | 96 |

### References

1. Rodier MH, Berthonneau J, Bourgoin A, Giraudeau G, Agius G, Burucoa C, et al. Seroprevalences of Toxoplasma, malaria, rubella, cytomegalovirus, HIV and treponemal infections among pregnant women in Cotonou, Republic of Benin. Acta tropica. 1995;59(4):271-7. Epub 1995/08/01. PubMed PMID: 8533662.

2. Dowdle WR, Ferrera W, De Salles Gomes LF, King D, Kourany M, Madalengoitia J, et al. WHO collaborative study on the sero-epidemiology of rubella in Caribbean and Middle and South American populations in 1968. Bull World Health Organ. 1970;42(3):419-22. Epub 1970/01/01. PubMed PMID: 5310208; PubMed Central PMCID: PMCPMC2427532.

3. Dutta SR, Atrash HK, Mathew L, Mathew PP, Mahmood RA. Seroepidemiology of rubella in Bahrain. Int J Epidemiol. 1985;14(4):618-23. Epub 1985/12/01. PubMed PMID: 4086149.

4. Rawls WE, Melnick JL, Bradstreet CM, Bailey M, Ferris AA, Lehmann NI, et al. WHO collaborative study on the sero-epidemiology of rubella. Bull World Health Organ. 1967;37(1):79-88. Epub 1967/01/01. PubMed PMID: 5300057; PubMed Central PMCID: PMC2554213.

5. Nessa A, Islam MN, Tabassum S, Munshi SU, Ahmed M, Karim R. Seroprevalence of rubella among urban and rural Bangladeshi women emphasises the need for rubella vaccination of pre-pubertal girls. Indian journal of medical microbiology. 2008;26(1):94-5. Epub 2008/01/30. PubMed PMID: 18227617.

6. Tahita MC, Hubschen JM, Tarnagda Z, Ernest D, Charpentier E, Kremer JR, et al. Rubella seroprevalence among pregnant women in Burkina Faso. BMC Infect Dis. 2013;13:164. doi: 10.1186/1471-2334-13-164. PubMed PMID: 23556510; PubMed Central PMCID: PMCPMC3623657.

7. Modarres S, Modarres S, Oskoii NN. The immunity of children and adult females to rubella virus infection in Tehran. Iranian journal of medical sciences. 1996;21:69-73.

8. Seth P, Manjunath N, Balaya S. Rubella infection: the Indian scene. Rev Infect Dis. 1985;7 Suppl 1:S64-7. Epub 1985/03/01. PubMed PMID: 4001736.

9. Mao B, Chheng K, Wannemuehler K, Vynnycky E, Buth S, Soeung SC, et al. Immunity to polio, measles and rubella in women of child-bearing age and estimated congenital rubella syndrome incidence, Cambodia, 2012. Epidemiol Infect. 2014:1-10. Epub 2014/11/07. doi: S0950268814002817 [pii] 10.1017/S0950268814002817. PubMed PMID: 25373419.

10. Yala F, Biendo M, Odongo I, Kounkou R. [Virological and bacteriological study of materno-fetal infections in Brazzaville]. Bulletin de la Societe de pathologie exotique (1990). 1991;84(5 Pt 5):627-34. Epub 1991/01/01. PubMed PMID: 1819414.
11. Pereira F, Uez O. Rubella antibodies in female applicants for premarital health certificates in Mar del Plata, Argentina. Bull Pan Am Health Organ. 1986;20(2):179-85. Epub 1986/01/01. PubMed PMID: 3768598.
12. El-Khateeb MS, Tarawneh MS, Hijazi S, Kahwaji L. Seroimmunity to rubella virus in Jordanians. Public Health. 1983;97(4):204-7. Epub 1983/07/01. PubMed PMID: 6622640.
13. Glikmann G, Petersen I, Mordhorst CH. Prevalence of IgG-antibodies to mumps and measles virus in non-vaccinated children. Dan Med Bull. 1988;35(2):185-7. Epub 1988/04/01. PubMed PMID: 3359817.
14. Wannian S. Rubella in the People's Republic of China. Reviews of Infectious Diseases. 1985;7(Supp 1):S72.
15. Vrinat M, Dutertre J, Helies H, Ropero P. [A serological survey of rubella among pregnant women in Abidjan (author's transl)]. Medecine tropicale : revue du Corps de sante colonial. 1978;38(1):53-7. Epub 1978/01/01. PubMed PMID: 214661.
16. Hathout H, Al-Nakib W, Lilley H, Abo-Ahmed HS, Nosseir AF. Seroepidemiology of rubella in Kuwait: an alternative vaccination policy. Int J Epidemiol. 1978;7(1):49-53. Epub 1978/03/01. PubMed PMID: 659049.
17. Edmunds WJ, Gay NJ, Kretzschmar M, Pebody RG, Wachmann H. The pre-vaccination epidemiology of measles, mumps and rubella in Europe: implications for modelling studies. Epidemiol Infect. 2000;125(3):635-50. Epub 2001/02/24. PubMed PMID: 11218214; PubMed Central PMCID: PMCPMC2869647.
18. Macnamara FN, Mitchell R, Miles JA. A study of immunity to rubella in villages in the Fiji islands using the haemagglutination inhibition test. J Hyg (Lond). 1973;71(4):825-31. Epub 1973/12/01. PubMed PMID: 4520516; PubMed Central PMCID: PMC2130425.
19. Ouattara SA, Brettes JP, Kodjo R, Penali K, Gershy-Damet G, Sangare A, et al. [Seroepidemiology of rubella in the Ivory Coast. Geographic distribution]. Bulletin de la Societe de pathologie exotique et de ses filiales. 1987;80(4):655-64. Epub 1987/01/01. PubMed PMID: 2830995.
20. Souza VA, Moraes JC, Sumita LM, Camargo MC, Fink MC, Hidalgo NT, et al. Prevalence of rubella antibodies in a non-immunized urban population, Sao Paulo, Brazil. The Division of Immunization, CVE. Revista do Instituto de Medicina Tropical de Sao Paulo. 1994;36(4):373-6. Epub 1994/07/01. PubMed PMID: 7732269.
21. Bedrossian NK, Matossian R. Is there a rubella problem in Lebanon? Lebanese Medical Journal. 1985;35(1):31-8.
22. Alleman MM, Wannemuehler KA, Hao L, Perelygina L, Icenogle JP, Vynnycky E, et al. Estimating the burden of rubella virus infection and congenital rubella syndrome through a rubella immunity assessment among pregnant women in the Democratic Republic of the Congo: Potential impact on vaccination policy. Vaccine. 2016;34(51):6502-11. doi: 10.1016/j.vaccine.2016.10.059. PubMed PMID: 27866768.
23. Reiche EM, Morimoto HK, Farias GN, Hisatsugu KR, Geller L, Gomes AC, et al. [Prevalence of American trypanosomiasis, syphilis, toxoplasmosis, rubella, hepatitis B, hepatitis C, human immunodeficiency virus infection, assayed through serological tests among pregnant patients, from 1996 to 1998, at the Regional University Hospital Norte do Parana]. Rev Soc Bras Med Trop. 2000;33(6):519-27. Epub 2001/02/15. doi: S0037-86822000000600002 [pii]. PubMed PMID: 11175581.
24. Nejmi S. [Immunologic survey of rubella in Moroccan women in the Rabat region (study of antibodies inhibiting hemagglutination in 548 serums)]. Maroc medical. 1972;52(559):420-5. Epub 1972/07/01. PubMed PMID: 4642407.

25. Morgan-Capner P, Wright J, Miller CL, Miller E. Surveillance of antibody to measles, mumps, and rubella by age. BMJ. 1988;297(6651):770-2. Epub 1988/09/24. PubMed PMID: 3142541; PubMed Central PMCID: PMC1834398.
26. Chakravarty MS, Gupta B, Das BC, Mukherjee MK, Mitra AC, Sarkar JK. Seroepidemiological study of rubella in Calcutta. The Indian journal of medical research. 1976;64(1):87-92. Epub 1976/01/01. PubMed PMID: 1270103.
27. Iqbal A, Bokhari S. Occurrence of rubella antibody IgG in the general population. Mother and Child. 1997;35(1):17-22.
28. Ukkonen P. Rubella immunity and morbidity: impact of different vaccination programs in Finland 1979-1992. Scand J Infect Dis. 1996;28(1):31-5. Epub 1996/01/01. PubMed PMID: 9122630.
29. Khare S, Banerjee K, Padubidri V, Rai A, Kumari S, Kumari S. Lowered immunity status of rubella virus infection in pregnant women. The Journal of communicable diseases. 1987;19(4):391-5. Epub 1987/12/01. PubMed PMID: 3507446.
30. Lam SK. The seroepidemiology of rubella in Kuala Lumpur, West Malaysia. Bull World Health Organ. 1972;47(1):127-9. Epub 1972/01/01. PubMed PMID: 4538899; PubMed Central PMCID: PMCPMC2480804.
31. Ahmed R, Hashmi K, Ullah SE, Khanum T, Rafia A. Study of Prevalence of Immune Status in Adult Females For Rubella Virus Infection. Pakistan Journal of Biological Sciences. 2006;9(5):816.
32. Khare S, Gupta HL, Banerjee K, Kumari S, Kumari S, Gupta HL. Seroimmunity to rubella virus infection in young adult females in Delhi. The Journal of communicable diseases. 1990;22(4):279-80. Epub 1990/12/01. PubMed PMID: 2098435.
33. Doraisingham S, Goh KT. The rubella immunity of women of child-bearing age in Singapore. Annals of the Academy of Medicine, Singapore. 1981;10(2):238-41. Epub 1981/04/01. PubMed PMID: 7332291.
34. Hossain A. Seroepidemiology of rubella in Saudi Arabia. Journal of tropical pediatrics. 1989;35(4):169-70. Epub 1989/08/01. PubMed PMID: 2810460.
35. Malakmadze N, Zimmerman LA, Uzicanin A, Shteinke L, Caceres VM, Kasymbekova K, et al. Development of a rubella vaccination strategy: contribution of a rubella susceptibility study of women of childbearing age in Kyrgyzstan, 2001. Clinical infectious diseases : an official publication of the Infectious Diseases Society of America. 2004;38(12):1780-3. Epub 2004/07/01. doi: 10.1086/421018. PubMed PMID: 15227627.
36. Brown DWJ, Cutts FT, Joseph A. An evaluation of complementary epidemiological methods in a defined population in Southern India for estimating the burden of Congenital Rubella Syndrome. 2004.
37. Miyakawa M, Yoshino H, Yoshida LM, Vynnycky E, Motomura H, Tho le H, et al. Seroprevalence of rubella in the cord blood of pregnant women and congenital rubella incidence in Nha Trang, Vietnam. Vaccine. 2014;32(10):1192-8. Epub 2013/09/12. doi: S0264-410X(13)01188-2 [pii] 10.1016/j.vaccine.2013.08.076. PubMed PMID: 24021315.
38. Sandow D, Okubagzhi GS, Arnold U, Denkmann N. Seroepidemiological study in rubella in pregnant women in Gondar Region, northern Ethiopia. Ethiopian medical journal. 1982;20(4):173-8. Epub 1982/10/01. PubMed PMID: 7140729.
39. Desinor OY, Anselme RJ, Laender F, Saint-Louis C, Bien-Aime JE. Seroprevalence of antibodies against rubella virus in pregnant women in Haiti. Rev Panam Salud Publica. 2004;15(3):147-50. Epub 2004/04/21. PubMed PMID: 15096284.
40. Saeed A, Abu-Shagra S, Al-Rasheed R. Congenital Rubella Syndrome - revisited (letter). Saudi Med J. 1993;14(25-26).
41. Dumitrescu R, Mateescu M, Gaicu N, Comanescu D. Evaluation of the anti-rubella immunity levels on a lot of 5,000 sera from women at procreative age, tested by HAI, in Romania. Archives roumaines de pathologie experimentales et de microbiologie. 1989;48(3):253-63. Epub 1989/07/01. PubMed PMID: 2519635.

42. Cutts FT, Abebe A, Messele T, Dejene A, Enquselassie F, Nigatu W, et al. Sero-epidemiology of rubella in the urban population of Addis Ababa, Ethiopia. Epidemiol Infect. 2000;124(3):467-79. Epub 2000/09/12. PubMed PMID: 10982071; PubMed Central PMCID: PMC2810933.
43. Nabli B. [Seroepidemiology of rubella in Tunisia]. Bull World Health Organ. 1970;42(6):891-6. Epub 1970/01/01. PubMed PMID: 5312251; PubMed Central PMCID: PMCPMC2427574.
44. Aksakal FN, Maral I, Cirak MY, Aygun R. Rubella seroprevalence among women of childbearing age residing in a rural region: is there a need for rubella vaccination in Turkey? Jpn J Infect Dis. 2007;60(4):157-60. Epub 2007/07/24. PubMed PMID: 17642522.
45. Mefane C. Rubella antibodies in 1737 girls and women in Gabon. Afrique Medicale. 1985;24(226):29-32. PubMed PMID: 19852020650.
46. Strauss J, Dobahi SS, Danes L, Kopecky K, Svandova E. Serological survey of rubella in Yemen in 1985. Journal of hygiene, epidemiology, microbiology, and immunology. 1989;33(2):163-7. Epub 1989/01/01. PubMed PMID: 2768819.
47. Pehlivan E, Karaoglu L, Ozen M, Gunes G, Tekerekoglu MS, Genc MF, et al. Rubella seroprevalence in an unvaccinated pregnant population in Malatya, Turkey. Public Health. 2007;121(6):462-8. Epub 2007/01/16. doi: S0033-3506(06)00327-1 [pii] 10.1016/j.puhe.2006.09.021. PubMed PMID: 17222875.
48. Upreti SR, Thapa K, Pradhan YV, Shakya G, Sapkota YD, Anand A, et al. Developing rubella vaccination policy in Nepal--results from rubella surveillance and seroprevalence and congenital rubella syndrome studies. J Infect Dis. 2011;204 Suppl 1:S433-8. Epub 2011/06/17. doi: 10.1093/infdis/jir078. PubMed PMID: 21666196.
49. Lawn JE, Reef S, Baffoe-Bonnie B, Adadevoh S, Caul EO, Griffin GE. Unseen blindness, unheard deafness, and unrecorded death and disability: congenital rubella in Kumasi, Ghana. American journal of public health. 2000;90(10):1555-61. Epub 2000/10/13. PubMed PMID: 11029988; PubMed Central PMCID: PMCPMC1446363.
50. Gutierrez Trujillo G, Munoz O, Tapia Conyer R, Bustamante Calvillo ME, Alvarez y Munoz MT, Guiscafre Gallardo JP, et al. [The seroepidemiology of rubella in Mexican women. A national probability survey]. Salud publica de Mexico. 1990;32(6):623-31. Epub 1990/11/01. PubMed PMID: 2089638.
51. Sallam TA, Al-Jaufy AY, Al-Shaibany KS, Ghauth AB, Best JM. Prevalence of antibodies to measles and rubella in Sana'a, Yemen. Vaccine. 2006;24(37-39):6304-8. Epub 2006/07/04. doi: 10.1016/j.vaccine.2006.05.083. PubMed PMID: 16815602.
52. Sasmaz T, Kurt AO, Ozturk C, Bugdayci R, Oner S. Rubella seroprevalence in women in the reproductive period, Mersin, Turkey. Vaccine. 2007;25(5):912-7. Epub 2006/10/20. doi: S0264-410X(06)01010-3 [pii] 10.1016/j.vaccine.2006.09.033. PubMed PMID: 17049680.
53. Desudchit P, Chatiyanonda K, Bhamornsathit S. Rubella antibody among Thai women of childbearing age. Southeast Asian J Trop Med Public Health. 1978;9(3):312-6. Epub 1978/09/01. PubMed PMID: 311950.
54. Kombich JM, PC; Borus, PK Seroprevalence of Natural Rubella Antibodies among Antenatal Attendees at Moi Teaching and Referral Hospital, Eldoret, Kenya. Journal of Immunological Techniques in Infectious Diseases. 2012;1(1).
55. Yamamoto L, Mejia E, Lopez RM, Gallardo E, Gomez B. Susceptibility to rubella infection in females at high risk. Immune protection associated to population density. Tropical and geographical medicine. 1995;47(6):235-8. Epub 1995/01/01. PubMed PMID: 8650731.
56. Cumberland P, Shulman CE, Maple PA, Bulmer JN, Dorman EK, Kawuondo K, et al. Maternal HIV infection and placental malaria reduce transplacental antibody transfer and tetanus antibody levels in newborns in Kenya. J Infect Dis. 2007;196(4):550-7. Epub 2007/07/13. doi: JID37995 [pii] 10.1086/519845. PubMed PMID: 17624840.
57. Scott S, Cumberland P, Shulman CE, Cousens S, Cohen BJ, Brown DW, et al. Neonatal measles immunity in rural Kenya: the influence of HIV and placental malaria infections on placental transfer of antibodies and levels of antibody in maternal and cord serum samples. J Infect Dis.

2005;191(11):1854-60. Epub 2005/05/05. doi: JID33850 [pii] 10.1086/429963. PubMed PMID: 15871118.
58. Dromigny JA, Pecarrere JL, Ollivier G, Leroy F, Zeller HG. [Seroprevalence of rubella in pregnant women at Antananarivo. Study of 853 sera at the Pasteur Institute in Madagascar]. Archives de l'Institut Pasteur de Madagascar. 1996;63(1-2):53-5. Epub 1996/01/01. PubMed PMID: 12463018.
59. Barreto J, Sacramento I, Robertson SE, Langa J, de Gourville E, Wolfson L, et al. Antenatal rubella serosurvey in Maputo, Mozambique. Trop Med Int Health. 2006;11(4):559-64. Epub 2006/03/24. doi: 10.1111/j.1365-3156.2006.01577.x. PubMed PMID: 16553940.
60. Odelola HA. Rubella haemagglutination inhibiting antibodies in females of child-bearing age in western Nigeria. Journal of hygiene, epidemiology, microbiology, and immunology. 1978;22(2):190-4. Epub 1978/01/01. PubMed PMID: 570582.
61. Bukbuk DN, el Nafaty AU, Obed JY. Prevalence of rubella-specific IgG antibody in non-immunized pregnant women in Maiduguri, north eastern Nigeria. Central European journal of public health. 2002;10(1-2):21-3. Epub 2002/07/05. PubMed PMID: 12096678.
62. Suarez-Ognio L, Adrianzen A, Ortiz A, Martinez C, Whittembury A, Cabezudo E, et al. A rubella serosurvey in postpartum women in the three regions of Peru. Rev Panam Salud Publica. 2007;22(2):110-7. Epub 2007/11/03. PubMed PMID: 17976277.
63. Amina MD, Oladapo S, Habib S, Adebola O, Bimbo K, Daniel A. Prevalence of rubella IgG antibodies among pregnant women in Zaria, Nigeria. International Health. 2010;2(2):156-9. PubMed PMID: 20103221973.
64. Pitts OM, Ravenel JM, Finklea JF. Rubella immunity in Trinidad. American journal of epidemiology. 1969;89(3):271-6. Epub 1969/03/01. PubMed PMID: 5773423.
65. Dromigny JA, Nabeth P, Perrier Gros Claude JD. Evaluation of the seroprevalence of rubella in the region of Dakar (Senegal). Trop Med Int Health. 2003;8(8):740-3. Epub 2003/07/19. PubMed PMID: 12869096.
66. Corcoran C, Hardie DR. Seroprevalence of rubella antibodies among antenatal patients in the Western Cape. South African medical journal = Suid-Afrikaanse tydskrif vir geneeskunde. 2005;95(9):688-90. Epub 2005/12/06. PubMed PMID: 16327929.
67. Mwambe B, Mirambo MM, Mshana SE, Massinde AN, Kidenya BR, Michael D, et al. Sero-positivity rate of rubella and associated factors among pregnant women attending antenatal care in Mwanza, Tanzania. BMC Pregnancy Childbirth. 2014;14:95. doi: 10.1186/1471-2393-14-95. PubMed PMID: 24589180; PubMed Central PMCID: PMCPMC3975942.
68. Watts T. Rubella antibodies in a sample of Lusaka mothers. Medical journal of Zambia. 1983;17(4):109-10. Epub 1983/10/01. PubMed PMID: 6680556.
69. United Nations, Department of Economic and Social Affairs, Population Division, World Population Prospects. https://population.un.org/wpp/Download/Archive/Standard/2017.
70. The World Bank Data Catalog, World Development Indicators. https://datacatalog.worldbank.org/dataset/world-development-indicators/resource/08feba-8a62-11e6-ae22-56b6b64900022017.
71. UNITED NATIONS DEVELOPMENT PROGRAMME Human Development Reports Table 2. Human Development Index Trends, 1990-2015 http://hdr.undp.org/en/composite/trends2016.
72. IMF DataMapper. https://www.imf.org/external/datamapper/PPPPC@WEO/OEMDC/ADVEC/WEOWORLD/.
73. University of Groningen Growth and Development Centre, Maddison Database 2010. https://www.rug.nl/ggdc/historicaldevelopment/maddison/data/md2010_vertical.xlsx.
74. Organisation for Economic Co-operation and Development, OECD.Stat, Health Care Utilisation: Consultations. https://stats.oecd.org/index.aspx?queryid=30161.
75. WHO Global Health Observatory data repository Hib (Hib3) Immunization coverage estimates by country. https://apps.who.int/gho/data/node.main.A829?lang=en.

76. WHO European Health Information Gateway Health for All explorer. https://gateway.euro.who.int/en/hfa-explorer/#CtIn99Lev6.
77. United Nations, Department of Economic and Social Affairs, Statistics Division. Demographic Statistics Database, Households by age and sex of reference person and by size of household. http://data.un.org/Data.aspx?d=POP&f=tableCode:50.
78. United Nations, Department of Economic and Social Affairs, Statistics Division, Demographic Statistics Database, Population in households by type of household, age and sex. http://data.un.org/Data.aspx?d=POP&f=tableCode:329.
